# Integrated molecular, clinical, and ontological analysis identifies overlooked disease relationships

**DOI:** 10.1101/214833

**Authors:** Winston A. Haynes, Rohit Vashisht, Francesco Vallania, Charles Liu, Gregory L. Gaskin, Erika Bongen, Shane Lofgren, Timothy E. Sweeney, Paul J. Utz, Nigam H. Shah, Purvesh Khatri

**Author notes:** Corresponding author *Email addresses* (Nigam H. Shah), (Purvesh Khatri).

## Abstract

Existing knowledge of human disease relationships is incomplete. To establish a comprehensive understanding of disease, we integrated transcriptome profiles of 41,000 human samples with clinical profiles of 2 million patients, across 89 diseases. Based on transcriptome data, autoimmune diseases clustered with their specific infectious triggers, and brain disorders clustered by disease class. Clinical profiles clustered diseases according to the similarity of their initial manifestation and later complications, identifying disease relationships absent in prior co-occurrence analyses. Our integrated analysis of transcriptome and clinical profiles identified overlooked, therapeutically actionable disease relationships, such as between myositis and interstitial cystitis. Our improved understanding of disease relationships will identify disease mechanisms, offer novel therapeutic targets, and create synergistic research opportunities.

## 1. Introduction

A thorough understanding of human disease relationships has potential to unlock disease mechanisms, identify drug targets, and create synergistic research opportunities.^1, 2^ However, existing knowledge of disease relationships is incomplete and flawed. Previous research exploring molecular disease relationships have used genetic (genome wide associations studies^3, 4, 5, 6, 7, 8, 9, 10^ or phenome wide association studies^11, 12, 13, 14, 15^) and transcriptomic^16, 17, 18, 19^ data to provide insight into shared architectures of disease. These disease relationships, particularly those based on transcriptomics data, have been limited to a small number of data sets (generating less reproducible findings^20^) and diseases.^16, 17, 18, 19^ Clinical disease relationship research has leveraged co-occurence of diseases in clinical records to establish comorbidity networks, patient strata, disease clusters, and temporal disease trajectories.^21, 22, 23, 24, 25, 26, 27, 28, 29, 30, 31, 32, 33, 34, 15^ However, these methods do not consider phenotypic similarity of disease. For example, diabetes mellitus patients are usually diagnosed with either type 1 or type 2 diabetes, but not both. But, both sets of patients may experience hypoglycemia and are at risk for similar complications, such as macular degeneration. Traditional co-occurence based approaches will miss such relationships. Finally, ontologies such as ICD-10 and disease ontology embody our knowledge about disease relationships from the perspectives of biomedical researchers and health care providers.^35, 36, 37, 38, 39^ These molecular, clinical, and ontological data provide complementary, yet incomplete, views of the science, practice, and knowledge of medicine, respectively.

To date, no study has combined molecular, clinical, and ontological evidence to define disease relationships. We hypothesized that combining these complementary views of disease similarity would yield novel insights into disease relationships. We integrated publicly-available transcriptome profiles of over 41,000 samples from 104 diseases,^40^ electronic health records of over 2 million patients, and ontologies declaring disease relationships. Unlike previous studies that used one transcriptome dataset per disease, we analyzed multiple transcriptome datasets per disease using MetaIntegrator to identify more robust disease signatures.^40, 41^ Unlike co-occurrence based approaches, we quantified similarity amongst diseases based on their clinical profiles that include the initial manifestations and later complications seen in that disease. By incorporating both ontological and drug indication data with molecular and clinical data, we identified overlooked and therapeutically promising disease relationships.

## 2. Results

### 2.1. Defining Molecular and Clinical Disease Profiles

We performed an integrated multi-cohort analysis using MetaIntegrator to calculate transcriptomic profiles for 104 diseases and conditions based on 41,388 samples from 619 studies^42^ to create a molecular profile for each disease. MetaIntegrator has been repeatedly used to identify reproducible gene signatures across a broad range of conditions including organ transplantation, autoimmune diseases, cancer, vaccination, neurodegenerative diseases, and infections.^43, 44, 45, 46, 47, 48, 49, 20, 50, 51^ From these molecular profiles (comprised of an effect size for each gene), we computed pairwise correlation of all 104 diseases [Figure 1:“Disease Molecular Correlation”, Figure S1b].

**Figure 1:**
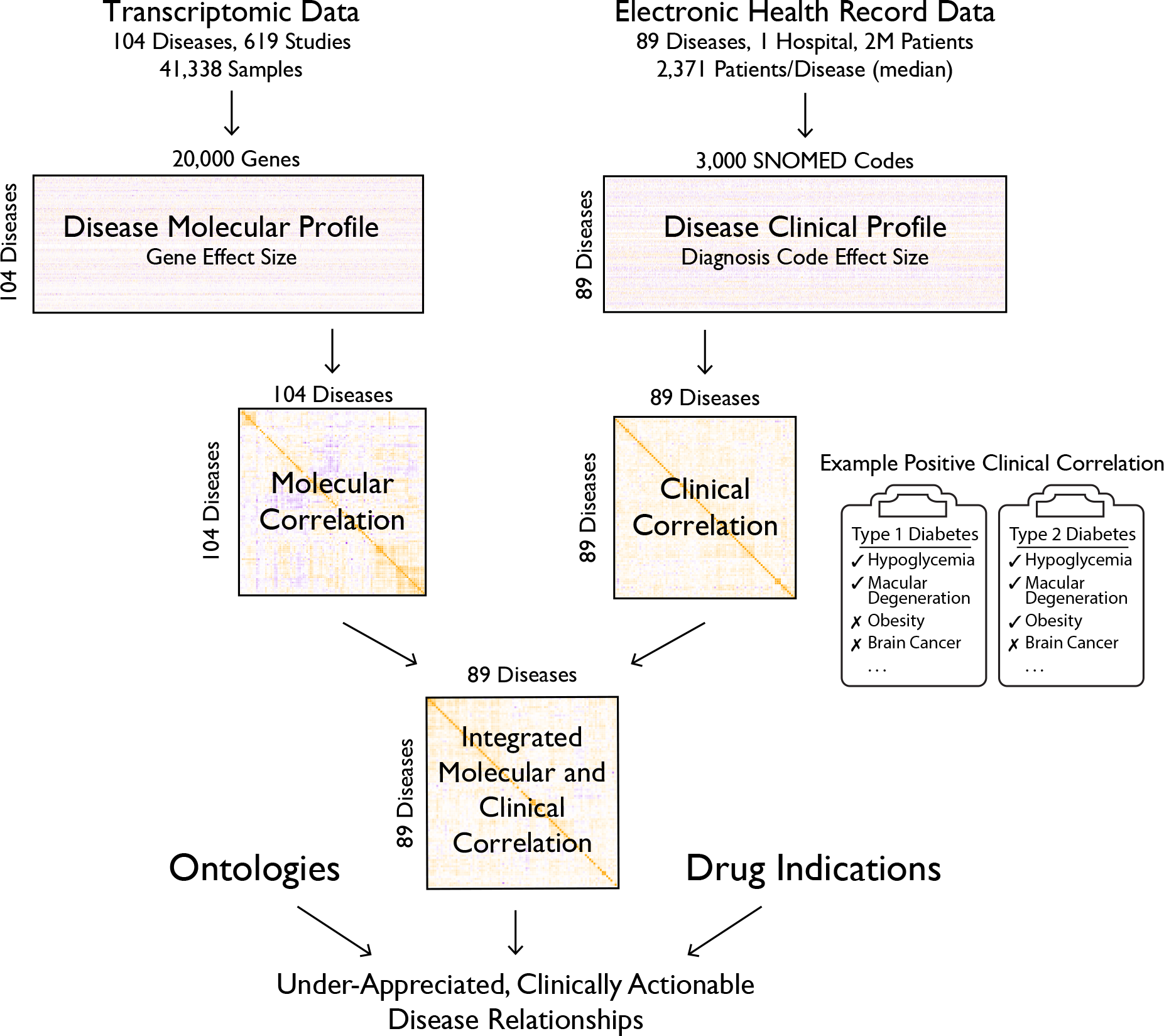
Integrated molecular, clinical, and ontological analysis for insight into disease similarity. Transcriptomic meta-analysis was performed to calculate effect sizes, a measure of differential expression, of genes across 104 diseases or conditions. In parallel, for 89 of those diseases, we analyzed electronic health records to define clinical profiles of diseases based on the association strength of a disease with all other diagnosis codes occurring in records of patients with that disease. From these effect sizes and clinical profiles, we calculated disease correlations. We performed a meta-analysis of disease correlations to calculate an integrated molecular and clinical disease correlation value. By integrating ontological and drug indication data, we identified overlooked, therapeutically actionable disease relationships.

We built clinical profiles for 89 diseases using the de-identified medical records of 2 million patients in the Stanford Clinical Data Warehouse^52^ [Figure 1]. The clinical profile of each disease *D* included the strength of association for 3,000 diagnosis codes with disease *D* compared to a matched cohort of patients without disease *D*. Unlike prior analyses that compared diseases based on co-occurrence, we quantified disease similarity based on the pairwise correlation of these clinical profiles.

### 2.2. Comparing Molecular and Clinical Disease Relationships

We performed hierarchical clustering of diseases based on molecular and clinical profiles to explore classes of related diseases [Figure 2]. Based on our molecular analysis, we highlight a few interesting clusters:

**Figure 2:**
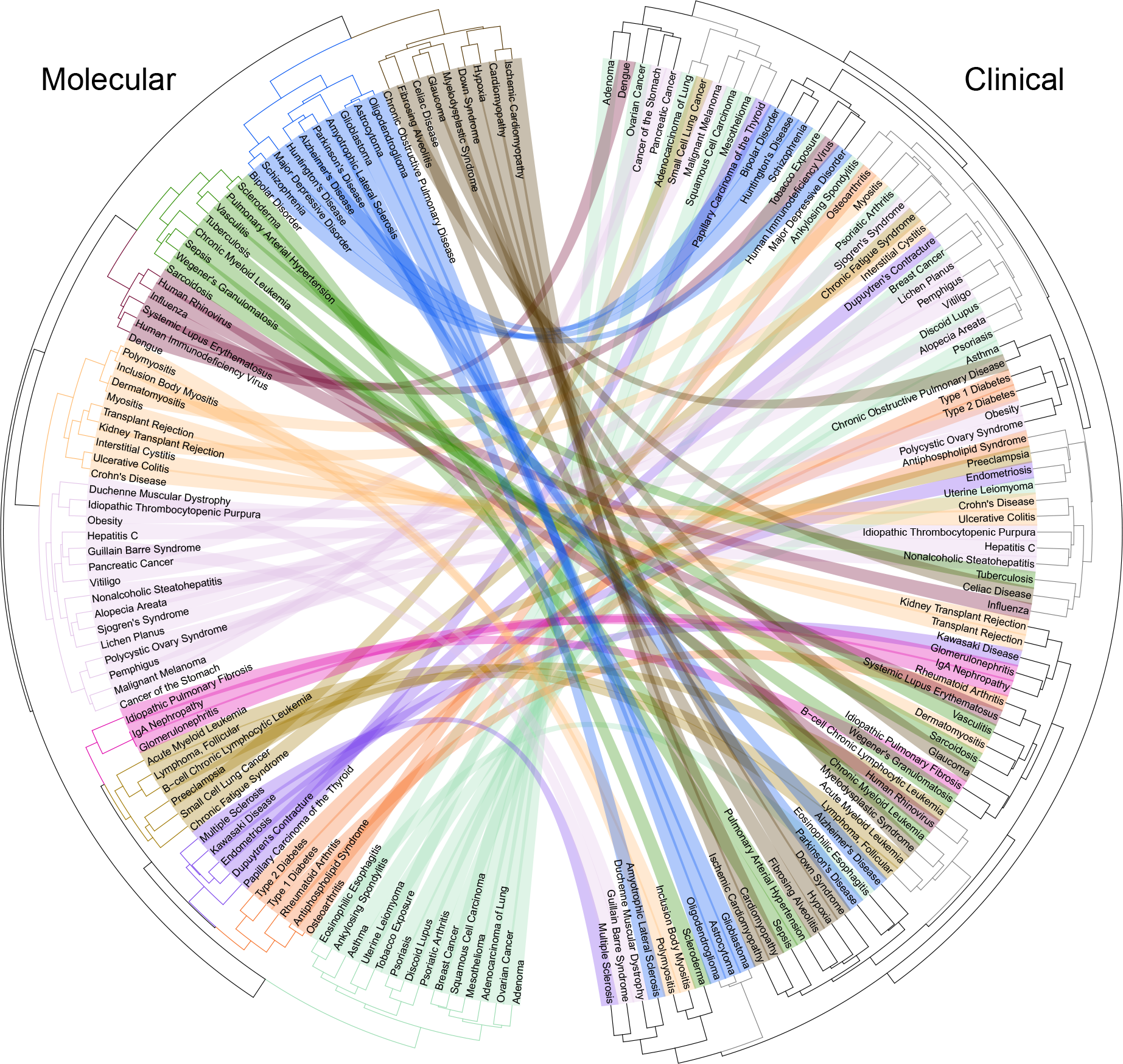
Comparing molecular and clinical perspectives of disease relationships. We created a dendrogram for molecular (left) and clinical (right) disease relationships based on pairwise disease correlations from our transcriptomic meta-analysis and electronic health record analysis, respectively. Arcs connect the same disease in the molecular and clinical dendrograms and are colored based on molecular groupings. See also Figure S1.

First, infectious diseases formed two distinct neighboring clusters, one containing all viral infections (maroon) and one containing all bacterial infections (green) [Figure 2]. These studies are concordant with our prior findings of a conserved viral signature^46^ and separate viral and bacterial signatures.^50^ The only non-infectious disease in the viral cluster was systemic lupus erythematosus, a complex autoimmune disease whose pathogenesis is strongly associated with Epstein Barr Virus infection.^53^ In the bacterial infection cluster: Wegener’s granulomatosis and sarcoidosis are autoimmune diseases that may be triggered by bacterial infection;^54, 55^ vasculitis is associated with both bacterial and viral infections;^56^ and pulmonary arterial hypertension is associated with both sepsis and HIV infection.^57, 58^ Thus, almost all diseases in these two clusters were either infectious or have been repeatedly associated with infection, which may explain the observed molecular relationships.

Second, a varied set of brain disorders formed a single cluster (blue) that further grouped into three distinct sub-clusters for neurodegenerative diseases, psychiatric disorders, and brain cancers [Figure 2]. Since it was possible that this cluster could have beenmay have been confounded by tissue-specific effects as we included transcriptome data from blood, solid tissue, and sorted cell types, we repeated our analysis using only blood transcriptomic profiles to avoid tissue bias. Despite using significantly less data, the transcriptomic analysis using solid tissues and blood datasets agreed significantly with the blood only datasets, in terms of both correlation and dendrogram similarity (p < 1e-4) [Figure S1a]. When using only blood transcriptomic data, Huntington’s disease, Alzheimer’s disease, and amyotrophic lateral sclerosis, the only neurological disorders with blood samples, were grouped in two neighboring clusters.

We compared dendrograms resulting from our clinical and molecular correlation analyses [Figure 2]. The clinical and molecular dendrograms demonstrated significant similarity (p < 1e-4). Despite this agreement, there were important differences between the disease relationships. In the clinical dendrogram, the brain cluster (blue), with the exception of Huntington’s disease and amyotrophic lateral sclerosis, split into three clusters which separately contained the brain cancers, psychiatric disorders, and neurodegenerative diseases. This split in neurological diseases reflects the differences in the clinical practice managing these diseases. On the other hand, the seemingly random dispersal of the bacterial (green) and viral (maroon) clusters was unexpected. It is possible that the wide variation in signs and symptoms of bacterial and viral infections (based on the site of the infection) leads to different clinical profiles for the infectious diseases.

### 2.3. Integrated Molecular and Clinical Disease Relationships

To combine the molecular and clinical perspectives, we performed a meta-analysis of the disease correlations [Figure 1:“Integrated Disease Correlation”, Figure S2b]. In the resulting hierarchical clustering, we observed that many of the disease groups were preserved: (1) bacterial (teal) and viral (blue) infection clusters contained many of the same members and were neighbors in the dendrogram; (2) brain diseases, except for amyotrophic lateral sclerosis, were entirely contained in two neighboring clusters (purple and grey) [Figure 3]. Further, we observed the emergence of a cluster of diseases that affects the muscles (maroon), including several types of myositis, Duchenne muscular dystrophy, and amyotrophic lateral sclerosis. Most of these clusters were reproducible at a p-value < 0.1 based on permutations of the molecular and clinical profiles data [Figure S2a]. These results demonstrate that the integrated analysis preserved meaningful groups identified from the molecular analysis and incorporated the clinical perspective.

**Figure 3:**
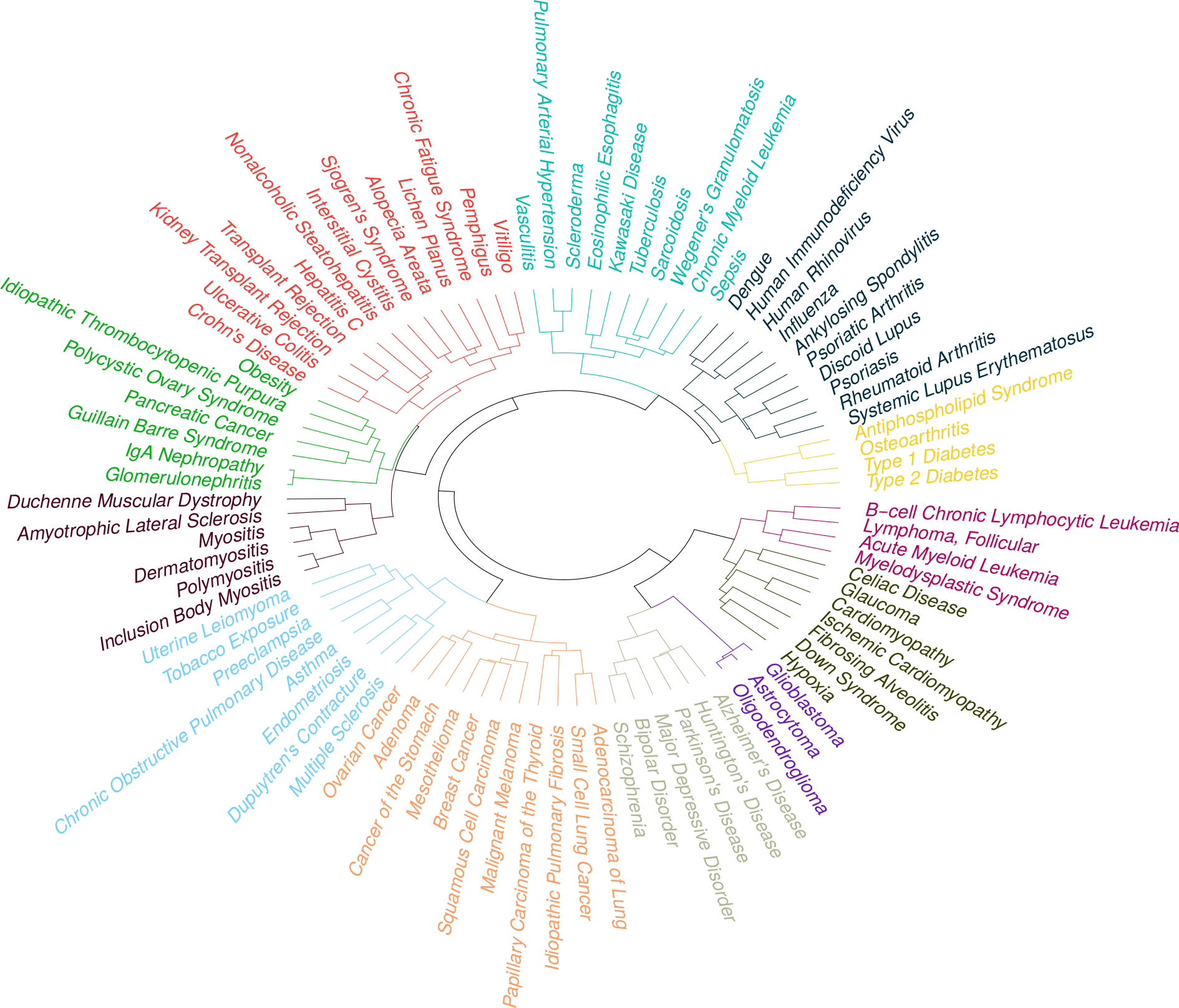
Integrated molecular and clinical disease relationships. Dendrogram resulting from meta-analysis of molecular and clinical disease correlations. See also Figure S2.

To illustrate the clinical utility of our integrated molecular and clinical analysis, we explored whether diseases with indications for the same drug cluster together in our analysis. If the clustering is clinically meaningful, then diseases treated with the same drug will be closer and more highly correlated than the background population. Indeed, we found that diseases treated with the same drugs were significantly closer and more similar to each other than other disease pairs (Wilcox test p-value < 1e-4 in all cases) [Figure S4]. Our results strongly suggest that disease pairs that are neighbors in the dendrogram or have highly positive correlation values should (or already do) respond to shared therapies.

### 2.4. Integration of Ontological Data to Identify Overlooked Molecular and Clincial Disease Relationships

When we compared disease relationships from our integrated molecular and clinical analysis with disease ontologies,^36, 35^ we observed significant similarities [Figure S2c]. In addition to the general agreement with prior knowledge, we found that disease pairs with the largest integrated correlation values were close to each other in the ontologies, including (1) astrocytoma and oligodendroglioma (brain cancers), (2) Huntington’s disease and Parkinson’s disease (neudegenerative conditions), and (3) malignant melanoma and squamous cell carcinoma (skin cancers) [Figure S3b]. Diseases with significantly negative integrated correlation values tended to be distant in the ontologies [Figure S3a].

We explored molecularly and clinically correlated disease pairs that were distant in ontologies and exhibited a disparity in the number of approved drugs [Figure 4]. Such disease pairs included (1) psoriasis and squamous cell carcinoma, (2) myositis and interstitial cystitis, (3) and ulcerative colitis and lichen planus. The majority of these top relationships have been co-mentioned in the same publication thirty times or fewer. Identifying such overlooked relationships has potential to benefit both disease research communities.^1, 2^

**Figure 4:**
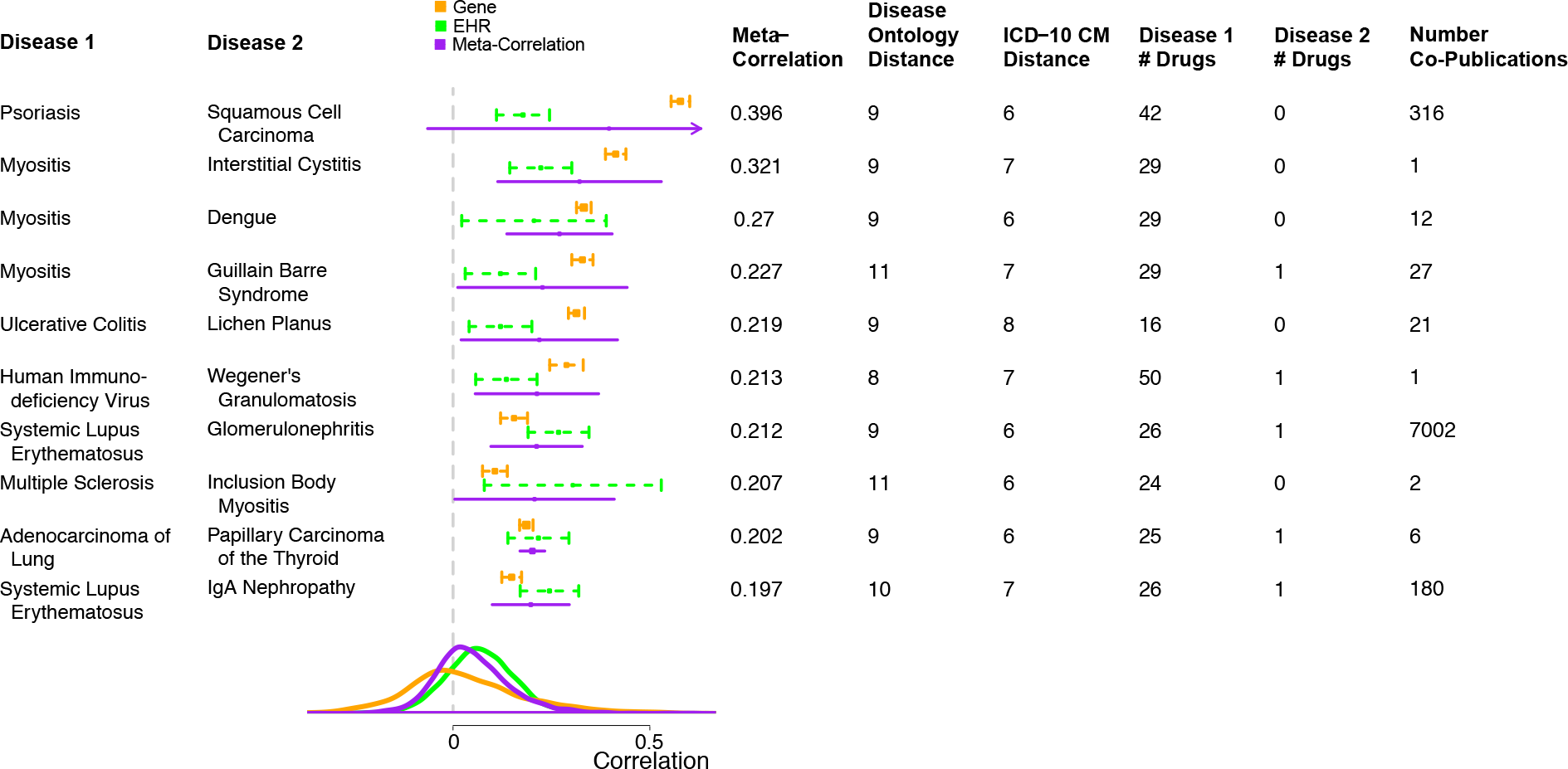
Overlooked, therapeutically promising disease correlations. The figure shows the top 10 disease pairs that were distant in both the Disease Ontology and ICD-10 CM, had a disparity in the number of available drugs, and had positive correlation based on molecular, clinical, and integrated analyses. The top 10 pairs are selected by the following criteria: at least 15 drugs in the MEDI database for disease 1, fewer than 3 drugs for disease 2, Disease Ontology distance greater than 7, ICD-10 CM distance greater than 5, and strictly positive 95% confidence intervals for both gene and EHR correlations. The density plot shows the background distribution of correlation values. See also Figure S3.

Consider the example of myositis and interstitial cystitis: myositis has been discussed in 17,503 PubMed publications and has 29 drugs in the MEDI database.^59^ In contrast, interstitial cystitis has been discussed in 1,824 publications and has no drugs in the MEDI database. Despite an integrated molecular and clinical correlation value of 0.32 between these diseases, the 27 million paper corpus of PubMed contains only two papers which mention both myositis and interstitial cystitis. We examined the genes with significant effect sizes in both diseases, including LOC100510692, LGALS9B, TMSB10, APOBEC3A, and SNORA14B [Figure S6]. With the exception of APOBEC3A, the other 4 top genes all have one or fewer Gene Ontology annotations, indicating that little is understood about these potentially important genes.^60, 61^ Gene Ontology enrichment analysis of all genes significant in both diseases at an FDR < 5% identified T cell apoptotic processes, microtubule organization, and cytokinesis as significant biological processes.^62^ We suggest that researchers pursue synergies between this and other significantly correlated disease pairs.

## 3. Discussion

We found that integration of molecular, clinical, and ontological data elucidated overlooked disease relationships. Identification of such unexpected, therapeutically actionable disease relationships can connect disparate research communities, improve understanding of disease causation, and generate novel therapeutic strategies.^1, 2^ In particular, we identify relationships like myositis and interstitial cystitis where deep knowledge about myositis may strengthen understanding and treatment of interstitial cystitis.

In the molecular analysis, we observed that infectious diseases separately clustered into bacterial and viral clusters alongside autoimmune diseases, whose onset may be triggered by those infections. We found that disease relationships based only on blood data were similar to relationships discovered when including all data (tissue, blood, and sorted cells), agreeing with prior findings that the effect of disease is stronger than the effect of tissue.^63^ This finding enables opportunities for studying diseases based on circulating blood when affected tissue such as brain, spleen, or pancreas may be technically challenging or dangerous to obtain.

In the clinical analysis, we provide the first quantification of disease similarity based on patient clinical profiles. Our approach captures the overall phenotypic similarity of diseases without the requirement that the same patients be diagnosed with both diseases. Although this work draws only on the Stanford electronic health record system, we are excited by future projects that improve reproducibility and generalizability by integrating multiple electronic health records systems.

Since the number of unique pairwise disease comparisons was too large to effectively visualize in a static graphic, we provide an interactive visualization of our data [http://metasignature.stanford.edu]. We additionally invite researchers to explore the underlying gene expression meta-analysis data to formulate their own hypotheses.

Although drug repositioning research focuses on dissimilarity of drug effects and disease state, we found that diseases with positive integrated correlation values were more likely to share drugs. Thus, we propose exploration of significantly correlated disease pairs for drug repositioning studies. By focusing on disease relationships, researchers can explore therapeutic options already available instead of limiting research to a single candidate molecule which may not be clinically effective.

By focusing on the science, practice, and knowledge of medicine, we increased our understanding of disease relationships. Our framework for integrating these diverse data types will be valuable for the new era of precision medicine, where opportunities for more deeply integrated analyses with both matched and timestamped clinical and molecular data will yield even more mechanistic and patient-specific insights. By increasing our knowledge of disease relatedness, we can better understand disease causation as well as provide ideas for therapeutic alternatives.

## 4. Materials and Methods

### 4.1. Gene expression data collection and meta-analysis

Gene expression meta-analysis data was compiled from the MetaSignature database.^40^ MetaSignature includes data from manual meta-analysis of over 41,000 samples, 619 studies, and 104 diseases and conditions [Supplemental Table S1]. Briefly, relevant data were downloaded from Gene Expression Omnibus and ArrayExpress.^64, 65^ Cases and controls were manually labeled for each disease and meta-analysis was performed using the MetaIntegrator package.^40^ We calculated the Hedges’ *g* summary effect size, standard error, and false discovery rate for every gene using the MetaIntegrator package.

For blood-only gene expression meta-analysis, we manually identified datasets that were derived from whole blood or peripheral blood mononuclear cells. We re-ran the MetaIntegrator analysis pipeline using only these datasets. Similarly, for the tissue and cell-type specific meta-analysis, we identified datasets derived from either solid tissue or sorted cell populations and ran MetaIntegrator using only these datasets. In total, 35 and 89 diseases had at least one dataset derived from blood and solid tissue/sorted cell, respectively.

### 4.2. Data collection for disease-gene publications, Single Nucleotide Polymorphism (SNP) data, and Gene Ontology annotations

We downloaded the number of publications for each disease-gene relationship from PubPular and HuGE Navigator for as many of the 104 diseases and conditions in MetaSignature as were present in the databases (102 in PubPular and 81 in HuGE).^66, 67, 68^ PubPular provided the top 261 gene associations, and HuGE provided all known associations. For all correlations, we only considered disease-gene associations with at least 10 publications to limit false positive associations.

We downloaded disease-SNP relationships, including gene mappings, odds ratios, and p-values, from the GWAS Catalog and HuGE Navigator for 61 and 54, respectively, of the 104 diseases in MetaSignature.^69, 70^ From Gene Ontology, we calculated the counts of non-Inferred from Electronic Annotation annotations for all the genes in the MetaSignature database.^60^ The Spearman rank correlation was used for all correlations.

Our plots show the top 10,0000 gene associations for each disease by effect size FDR rank. Correlation calculations do not include a similar limit.

### 4.3. EHR data collection and analysis

Patient level data were obtained from Stanford Clinical Data Warehouse (CDW), which represents electronic medical records (EMRs) of patients from both Lucile Packard Children’s Hospital and Stanford Hospitals and clinics.^52^ The CDW contains data on about 2 million adult and pediatric patients consisting of demographic information, 25 million clinical encounters, 48 million inpatient and outpatient ICD9-CM diagnosis codes, 157 million laboratory test results along with various types of pharmacy orders, clinical text, surgical reports and radiology reports. The data were converted into OMOP common data model v5.^71^ The common data model allows consistent representation of the data in using common terminologies such as SNOMED-CT, RxNorms, and LOINC. Of the 104 diseases from the gene expression meta-analysis, we focused on 89, which clearly mapped to diagnosis codes and had at least 100 patients in STRIDE.

The clinical profile of a disease D is a vector of length i, where each feature is another disease Di and the values are the strength of association between D and Di obtained from a matched cohort of patient with and without disease D. We calculated clinical profiles of diseases as follows: The ICD-9 CM diagnosis codes corresponding to each of the 89 diseases were manually curated using literature searches as well as prior published studies on those diseases. For example, for Type-2 diabetes we selected the codes 250.00, 250.02, 250.10, 250.12, 250.20, 250.22, 250.30, 250.32, 250.40, 250.42, 250.50, 250.52, 250.60, 250.62, 250.70, 250.72, 250.80, 250.82, 250.90, 250.92. Each code was then mapped to its corresponding SNOMED-CT code based on the mapping available from Unified Medical Language System.^72^ Cohorts of patients with and without disease were identified based on the presence or absence of the SNOMED-CT codes in a patient medical record [Figure S5].

The first occurrence of a SNOMED-CT code in the longitudinal record of a patient defined an index date. A patient was included in the cohort if at least 90 days of medical data were available. We also included demographic information such as age and gender. The control group of patients was identified by 1:1 matching of patients using a propensity score model adjusting for age, gender, and total length of record [Figure S5b]. The set of patients with the disease, and their matching controls, were then used to build a time agnostic, binary patient feature matrix capturing the occurrence of all other diseases that are found in the medical records of the individuals with the disease or in the records of the matching controls [Figure S5d]. For each disease, we obtained a matrix in which patients were in rows, and columns were binary indicators for the presence or absence of SNOMED-CT codes in the medical records of a given patient. To obtain a clinical profile of the disease from this matrix, we fit a ridge regression with presence of the disease of interest as the outcome. The resulting clinical profile of a disease was comprised of coefficients for each of the binary indicators (i.e. the SNOMED-CT codes). The exponent of the coefficient represents the effect size of association of a specific indicator in the profile of the given disease. Thus the resulting clinical profile of a disease D is a vector of length i, where each feature is another disease Di and the values are the strength of association between D and Di.

We used 10-fold cross validation while fitting the regression, and coefficients were selected for most regularized lambda. For each cohort, we bootstrapped the ridge regression for the fixed value of lambda (obtained from 10-fold cross validation) to compute the variance associated with the coefficients. Clinical profiles were constructed for each of the 89 diseases under study, and then were used to calculate pair-wise correlations between each disease.

Earlier approaches to infer associations between diseases relied on co-frequency,^21, 7^ which does not control for confounding comorbidities that might drive an observed association. For example, consider Diabetes Mellitus Type 2 (T2DM), Retinopathy and, Renal failure. An association between Retinopathy and Renal failure based on co-frequency will not account for the fact that an observed association between the two could be due to the high association between T2DM and Retinopathy as well as the strong association between T2DM and Renal failure. Using a regression “adjusts” for such effects. To illustrate the effect of such adjustment, we computed both the un-adjusted and adjusted odds ratios for all the disease pairs. 2351 disease pairs lost their positive association based on co-frequency, thus highlighting the importance of adjusting odds ratios when computing associations.

The R packages matching^73^ and glmnet^74^ were utilized to perform patient-level matching and ridge regression respectively, while custom SQL queries were written to extract patient-level data from the CDW.

### 4.4. Correlation calculation

We followed the same process for gene expression and electronic health record data, so we generically refer to each of these matrices as a matrix of features. For each pair of diseases, we calculated the Spearman correlation of all features that were significant at a FDR of less than 5% in either disease. We calculated confidence intervals and p-values for these correlations. We adjusted p-values using the Bonferroni correction.

### 4.5. Integrated molecular and clinical correlation analysis

We used meta-analysis of correlation values to combine disease correlations from clinical and molecular datasets. We converted correlation values from gene expression and EHR to Fisher’s z statistic and applied an inverse variance, random effects meta-analysis model to calculate our integrated correlation value.^75^ To calculate empirical p-values for the correlations, we performed 10,000 bootstrap resamplings of the gene expression and EHR data.

### 4.6. Clustering calculation

We converted correlation values to a distance as: 1 – cor. We performed hierarchical clustering of correlation distances using Ward’s clustering criterion.^76, 77^ To assess the robustness of our clustering, we adapted the pvclust R package and performed 10,000 bootstrap resamplings of the terms used for calculating correlation.^78^ For visualization, we leveraged the circlize R package.^79^

### 4.7. Ontology analysis

We downloaded OWL formatted data from the Disease Ontology and ICD-10 CM.^35, 36^ We mapped all MetaSignature diseases and conditions to SNOMED CUI, which map to 104 diseases in ICD-10 CM and 97 diseases in the Disease Ontology. Using the rdflib library in Python, we calculated the number of steps in the shortest path between pairs of terms by traversing "subClassOf" relationships. From these distances, we calculated a hierarchical clustering of terms using Ward’s clustering criterion.^76, 77^ The number of co-publications was determined through a PubMed search using both disease names connected with the ‘AND’ operator.

### 4.8. Dendrogram similarity

To numerically compare pairs of dendrograms, we pruned them to include only diseases present in both dendrograms, and calculated both the Pearson correlation of the underlying correlation matrices and the correlation of the cophenetic distance matrices.^80^ We used the R package dendextend for calculating the correlation of the cophenetic distance matrices.^81^ To empirically calculate the significance of both of these correlations, we performed 10,000 random permutations of the dendrogram and correlation matrix labels. For visualization, we used chord diagrams from the R package circlize and tanglegrams from the dendextend package.^81, 79^

### 4.9. Drug analysis

We downloaded drug indication data from Therapeutics Target DB and MEDI.^82, 59^ We limited our analysis to drugs with less than ten distinct disease indications matching to the Metasignature database. From Therapeutics Target DB and MEDI we identified 67 and 153 pairs of diseases from MetaSignature with indications for the same drug. We compared the similarity of diseases which share drugs in these databases to all other disease pairs.

### 4.10. Example disease pair

We explored the relationship of myositis and interstitial cystitis. We identified all genes with an FDR of less than 5% in both diseases. We highlighted those genes with an effect size of greater than 1 or less than 1 in both diseases. For the significant genes, we performed Gene Ontology enrichment analysis using the PANTHER Gene Ontology enrichment analysis tool (Analysis Type: PANTHER Overrepresentation Test (release 20170413), Annotation Version and Release Date: GO Ontology database Released 2017-04-24, Reference List: Homo sapiens (all genes in database), Annotation Data Set: GO biological process complete).^62^

## 5. Author Contributions

Conceptualization, W.A.H., R.V., P.J.U., N.H.S., and P.K; Methodology, W.A.H., R.V., N.H.S., and P.K; Software, W.A.H. and R.V.; Investigation, W.A.H. and R.V.; Data Curation, W.A.H., R.V., F.V., C.L, G.L.G, E.B., S.L., T.E.S.; Writing-Original Draft, W.A.H., R.V., P.J.U. N.H.S., and P.K.; Writing-Reviewing and Editing, W.A.H., R.V., F.V., C.L, G.L.G, E.B., S.L., T.E.S., P.J.U. N.H.S., and P.K.; Visualization, W.H.; Funding Acquisition, P.J.U., N.H.S., and P.K.

## 6. Acknowledgements

We thank Alex Schrenchuk for computer support. WAH is funded by the National Science Foundation Graduate Research Fellowship under Grant No. DGE-114747. FV is funded by the National Institute of Health K12 Career Award 5K12HL120001-02. EB is funded by Gabilan Fellowship. PK is funded by the National Institute of Allergy and Infectious Diseases grants 1U19AI109662, U19AI057229, U54I117925, and U01AI089859. PJU is the recipient of a gift from Gustav Floren Family Trust, and a gift from The Ben May Charitable Trust of Mobile, Alabama. PJU is supported by NIH grants NIAID U19-AI1110491, Stanford Autoimmunity Center of Excellence (ACE); NIAID 1 UM1AI110498-01, ACE Collaborative Project; 1 U19-AI090019; amd NIAID 1 R01 AI125197-01. NHS is supported in part by NLM grant R01 LM011369, NIGMS grant R01 GM101430, and a research award from Janssen R&D LLC.

## 8. Supplemental Information

**Table S1:** Sample size and number of studies for all 104 diseases in MetaSignature.

| Disease Name | Sample Size | Number of Studies |
| --- | --- | --- |
| Acute Myeloid Leukemia | 64 | 2 |
| Acute Promyelocytic Leukemia | 10 | 1 |
| Adenocarcinoma of Lung | 1524 | 17 |
| Adenoma | 38 | 2 |
| Alopecia Areata | 17 | 2 |
| Alzheimer's Disease | 997 | 9 |
| Amyotrophic Lateral Sclerosis | 38 | 2 |
| Ankylosing Spondylitis | 56 | 3 |
| Antiphospholipid Syndrome | 6 | 1 |
| Asthma | 130 | 2 |
| Astrocytoma | 83 | 2 |
| B-cell Chronic Lymphocytic Leukemia | 124 | 3 |
| Bacterial Infection | 82 | 3 |
| Bipolar Disorder | 219 | 6 |
| Breast Cancer | 1840 | 15 |
| Cancer of the Stomach | 43 | 2 |
| Cardiomyopathy | 706 | 13 |
| Celiac Disease | 1372 | 10 |
| Chronic Fatigue Syndrome | 148 | 5 |
| Chronic Myeloid Leukemia | 56 | 2 |
| Chronic Obstructive Pulmonary Disease | 58 | 4 |
| Crohn's Disease | 160 | 5 |
| Dengue | 687 | 4 |
| Dermatomyositis | 294 | 11 |
| Discoid Lupus | 38 | 1 |
| Down Syndrome | 25 | 2 |
| Duchenne Muscular Dystrophy | 137 | 3 |
| Dupuytren's Contracture | 34 | 4 |
| Endometriosis | 241 | 9 |
| Eosinophilic Esophagitis | 10 | 1 |
| Familial Combined Hyperlipidemia | 33 | 2 |
| Fibrosing Alveolitis | 6 | 1 |
| Glaucoma | 78 | 2 |
| Glioblastoma | 83 | 2 |
| Glomerulonephritis | 102 | 4 |
| Guillain Barre Syndrome | 14 | 1 |
| Hepatitis C | 106 | 5 |
| Human Immunodeficiency Virus | 133 | 3 |
| Human Rhinovirus | 140 | 3 |
| Huntington's Disease | 587 | 6 |
| Hypoxia | 10 | 2 |
| Idiopathic Fibrosing Alveolitis | 294 | 9 |
| Idiopathic Pulmonary Fibrosis | 340 | 8 |
| Idiopathic Thrombocytopenic Purpura | 4 | 1 |
| IgA Nephropathy | 195 | 7 |
| Inclusion Body Myositis | 17 | 1 |
| Influenza | 1346 | 13 |
| Interstitial Cystitis | 57 | 4 |
| Ischemic Cardiomyopathy | 38 | 2 |
| Kawasaki Disease | 89 | 2 |
| Kidney Transplant Rejection | 1486 | 11 |
| Lichen Planus | 47 | 3 |
| Major Depressive Disorder | 62 | 2 |
| Malignant Melanoma | 61 | 2 |
| Mesothelioma | 108 | 4 |
| Meta-Bacterial | 726 | 8 |
| Meta-Virus | 437 | 9 |
| Multiple Sclerosis | 1668 | 19 |
| Myelodysplastic Syndrome | 89 | 2 |
| Myositis | 273 | 11 |
| Narcolepsy | 20 | 1 |
| Non-small Cell Lung Cancer | 1339 | 9 |
| Nonalcoholic Steatohepatitis | 92 | 3 |
| Obesity | 281 | 4 |
| Oligodendroglioma | 83 | 2 |
| Osteoarthritis | 46 | 2 |
| Ovarian Cancer | 386 | 7 |
| Pancreatic Cancer | 294 | 9 |
| Papillary Carcinoma of the Thyroid | 32 | 2 |
| Parkinson's Disease | 236 | 11 |
| Pemphigus | 24 | 1 |
| Polycystic Ovary Syndrome | 70 | 3 |
| Polymyositis | 17 | 1 |
| Preeclampsia | 346 | 12 |
| Pregnancy | 171 | 3 |
| Primary Open Angle Glaucoma | 9 | 2 |
| Psoriasis | 488 | 12 |
| Psoriatic Arthritis | 38 | 2 |
| Pulmonary Arterial Hypertension | 227 | 4 |
| Respiratory Syncytial Virus | 251 | 6 |
| Rheumatoid Arthritis | 680 | 20 |
| Sarcoidosis | 487 | 10 |
| Schizophrenia | 305 | 8 |
| Scleroderma | 562 | 14 |
| Senescence | 101 | 5 |
| Sepsis | 788 | 9 |
| Sjogren's syndrome | 35 | 1 |
| Small Cell Lung Cancer | 581 | 6 |
| Squamous Cell Carcinoma | 36 | 2 |
| Systemic Lupus Erythematosus | 3748 | 21 |
| Tobacco Exposure | 603 | 8 |
| Transplant Rejection | 251 | 8 |
| Type 1 Diabetes | 1196 | 19 |
| Type 2 Diabetes | 122 | 2 |
| Ulcerative Colitis | 467 | 20 |
| Uterine Leiomyoma | 38 | 3 |
| Vasculitis | 29 | 1 |
| Vitiligo | 10 | 1 |
| Wegener's Granulomatosis | 127 | 1 |

**Table S2:** Number of patients for each of the 89 diseases we examined from the Stanford EHR.

| Disease Name | Number of Patients |
| --- | --- |
| Acute Myeloid Leukemia | 3172 |
| Adenocarcinoma of Lung | 6214 |
| Adenoma | 3785 |
| Alopecia areata | 743 |
| Alzheimers disease | 2339 |
| Amyotrophic Lateral Sclerosis | 379 |
| Ankylosing spondylitis | 842 |
| Antiphospholipid syndrome | 1446 |
| Asthma | 38580 |
| Astrocytoma | 2565 |
| B-cell Chronic Lymphocytic Leukemia | 4263 |
| Bipolar Disorder | 5934 |
| Breast cancer | 12718 |
| Cancer of the Stomach | 1850 |
| Cardiomyopathy | 11448 |
| Celiac disease | 1412 |
| Chronic fatigue syndrome | 1605 |
| Chronic myeloid leukemia | 1135 |
| Chronic obstructive pulmonary disease | 55856 |
| Crohns disease | 2996 |
| Dengue | 29 |
| Dermatomyositis | 506 |
| Discoid lupus | 524 |
| Down syndrome | 987 |
| Duchenne Muscular Dystrophy | 436 |
| Dupuytren's contracture | 1441 |
| Endometriosis | 7435 |
| Eosinophilic gastritis | 569 |
| Fibrosing alveolitis | 928 |
| Glaucoma | 12057 |
| Glioblastoma | 2565 |
| Glomerulonephritis | 2091 |
| Guillain Barre Syndrome | 562 |
| Hepatitis C | 9644 |
| Human immunodeficiency virus | 1646 |
| Human Rhinovirus | 144 |
| Huntingtons Disease | 65 |
| Hypoxia | 5711 |
| Idiopathic Thrombocytopenic Purpura | 651 |
| IgA nephropathy | 4275 |
| Inclusion body myositis | 26 |
| Influenza | 2246 |
| Interstitial Cystitis | 2000 |
| Ischemic Cardiomyopathy | 5714 |
| Kawasaki Disease | 267 |
| Kidney Transplant Rejection | 1462 |
| Lichen planus | 770 |
| Lymphoma Follicular | 1814 |
| Major depressive disorder | 8571 |
| Malignant melanoma | 3015 |
| Mesothelioma | 6965 |
| Multiple sclerosis | 2564 |
| Myelodysplastic Syndrome | 1454 |
| Myositis | 16397 |
| Nonalcoholic Steatohepatitis | 22142 |
| Obesity | 33551 |
| Oligodendroglioma | 2565 |
| Osteoarthritis | 37864 |
| Ovarian Cancer | 2758 |
| Pancreatic cancer | 2929 |
| Papillary Carcinoma of the Thyroid | 3214 |
| Parkinsons disease | 4055 |
| Pemphigus | 262 |
| Polycystic ovary syndrome | 2140 |
| Polymyositis | 240 |
| Preeclampsia | 2371 |
| Psoriasis | 4623 |
| Psoriatic Arthritis | 1060 |
| Pulmonary arterial hypertension | 7467 |
| Pulmonary fibrosis | 6524 |
| Rheumatoid Arthritis | 1031 |
| Sarcoidosis | 1477 |
| Schizophrenia | 2613 |
| Scleroderma | 1073 |
| Sepsis | 9127 |
| Sjogrens syndrome | 1141 |
| Small Cell Lung Cancer | 6214 |
| Squamous Cell Carcinoma | 1668 |
| Systemic Lupus Erythematosus | 3869 |
| Tobacco Exposure | 19568 |
| Transplant Rejection | 3528 |
| Tuberculosis | 1513 |
| Type 1 Diabetes | 7882 |
| Type 2 Diabetes | 45544 |
| Ulcerative colitis | 3573 |
| Uterine Leiomyoma | 13325 |
| Vasculitis | 1255 |
| Vitiligo | 953 |
| Wegener's granulomatosis | 300 |

**Figure S1:**
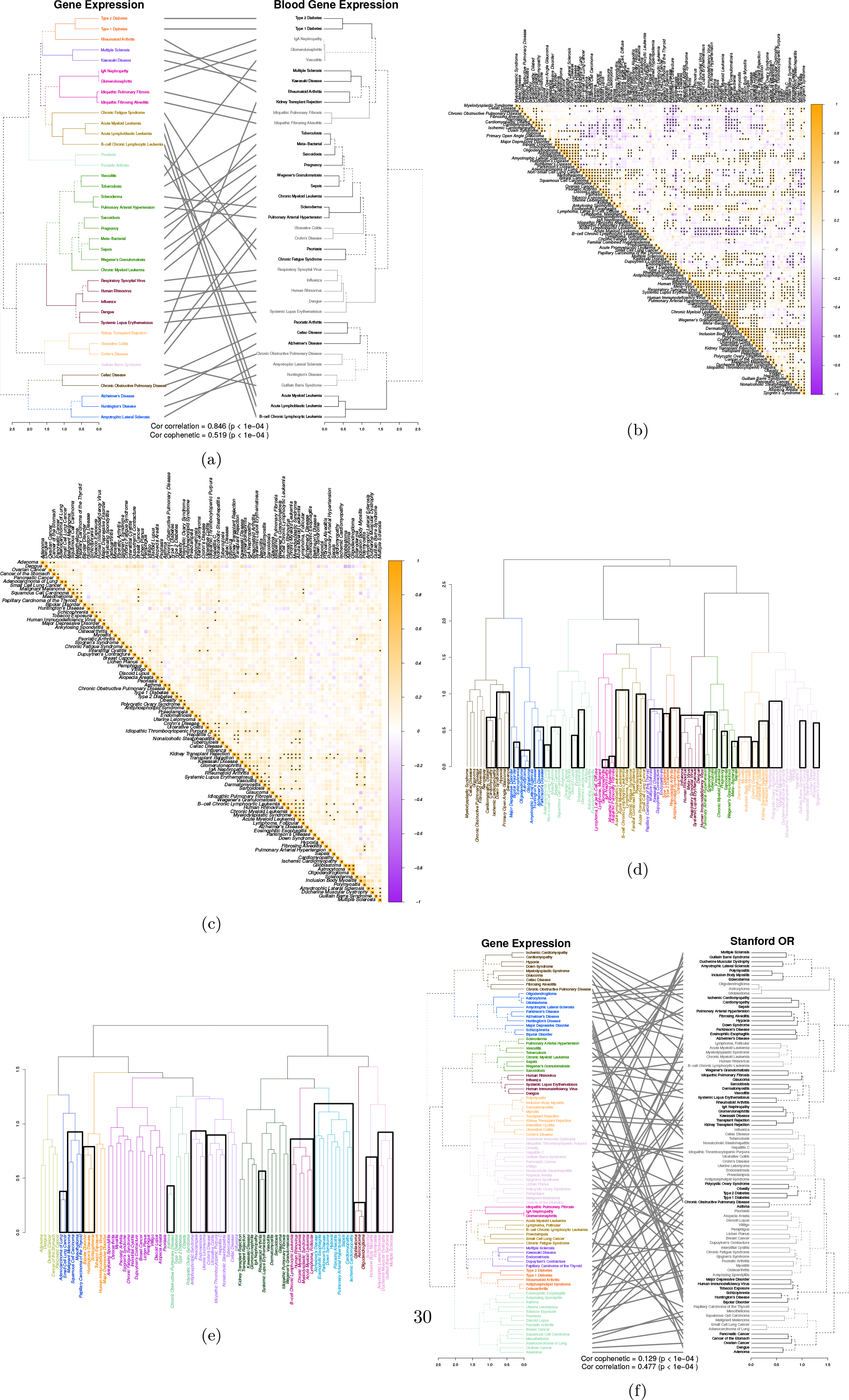
Related to Figure 2. (a) Gene expression clustering based on all samples vs. blood only samples. (b) Pairwise correlations of diseases based on gene expression meta-analysis. Stars indicate Bonferroni corrected p-values < 1e-10. (c) Pairwise correlations of diseases based on Stanford electronic health records. Stars indicate Bonferroni corrected p-values < 1e-5. (d) Bootstrap reproducibility of gene expression dendrogram. Outlined branches are reproducible at approximately unbiased p-value < 0.1. (e) Bootstrap reproducibility of electronic health record dendrogram. Outlined branches are reproducible at approximately unbiased p-value < 0.1. (f) Clustering based on gene expression vs. electronic health records.

**Figure S2:**
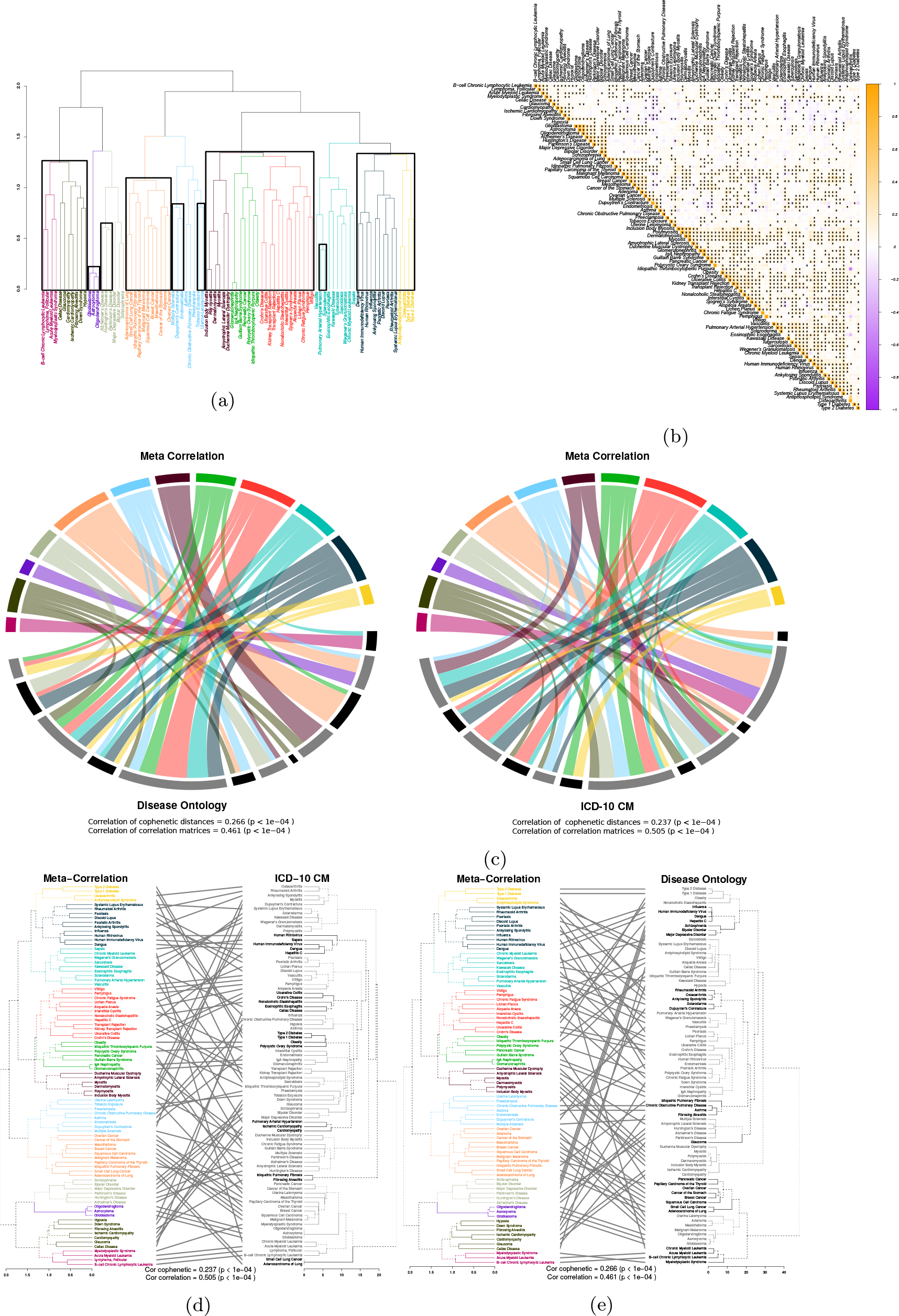
Related to Figure 3. (a) Bootstrap reproducibility of integrated molecular and clinical correlation dendrogram. Outlined branches are reproducible at approximately unbiased p-value < 0.1. (b) Pairwise correlations of diseases based on integrated molecular and clinical analysis. Stars indicate bootstrap p-values < 1e-3. (c) Comparison of the clustering from the meta-correlation analysis to the Disease Ontology and ICD10 Clinical Modification clusterings, respectively.^36, 35^ Colored arcs correspond to the clusters from Figure 3. Flows between arcs represent correspondence of disease cluster members. (d) Clustering based on meta-correlation vs. ICD-10 Clinical Modification. (e) Clustering based on metacorrelation vs. Disease Ontology.

**Figure S3:**
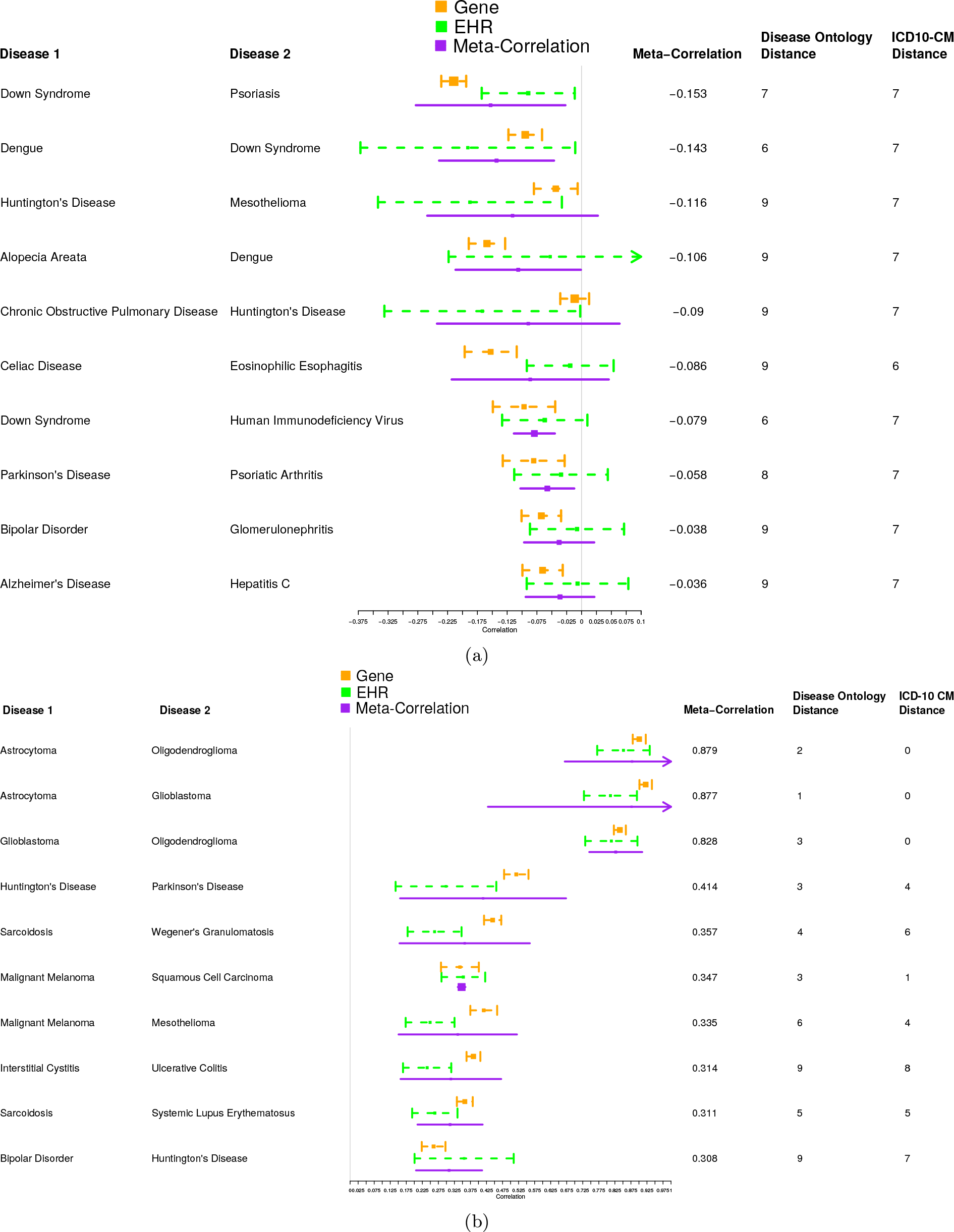
Related to Figure 4. (a) Meta-correlations with largest negative values. Displayed meta-correlations had negative meta-correlation values, gene expression and EHR correlations with the same directionality, and mappings to the Disease Ontology. The top 10 pairs of diseases in terms of meta-correlation that passed these criterion are displayed. (b) Sampling of significant pairwise meta-correlations. Displayed meta-correlations had a p-value < 1e-3, gene expression and EHR correlations with the same directionality, and mappings to the Disease Ontology. The top 10 pairs of diseases in terms of meta-correlation that passed these criterion are displayed.

**Figure S4:**
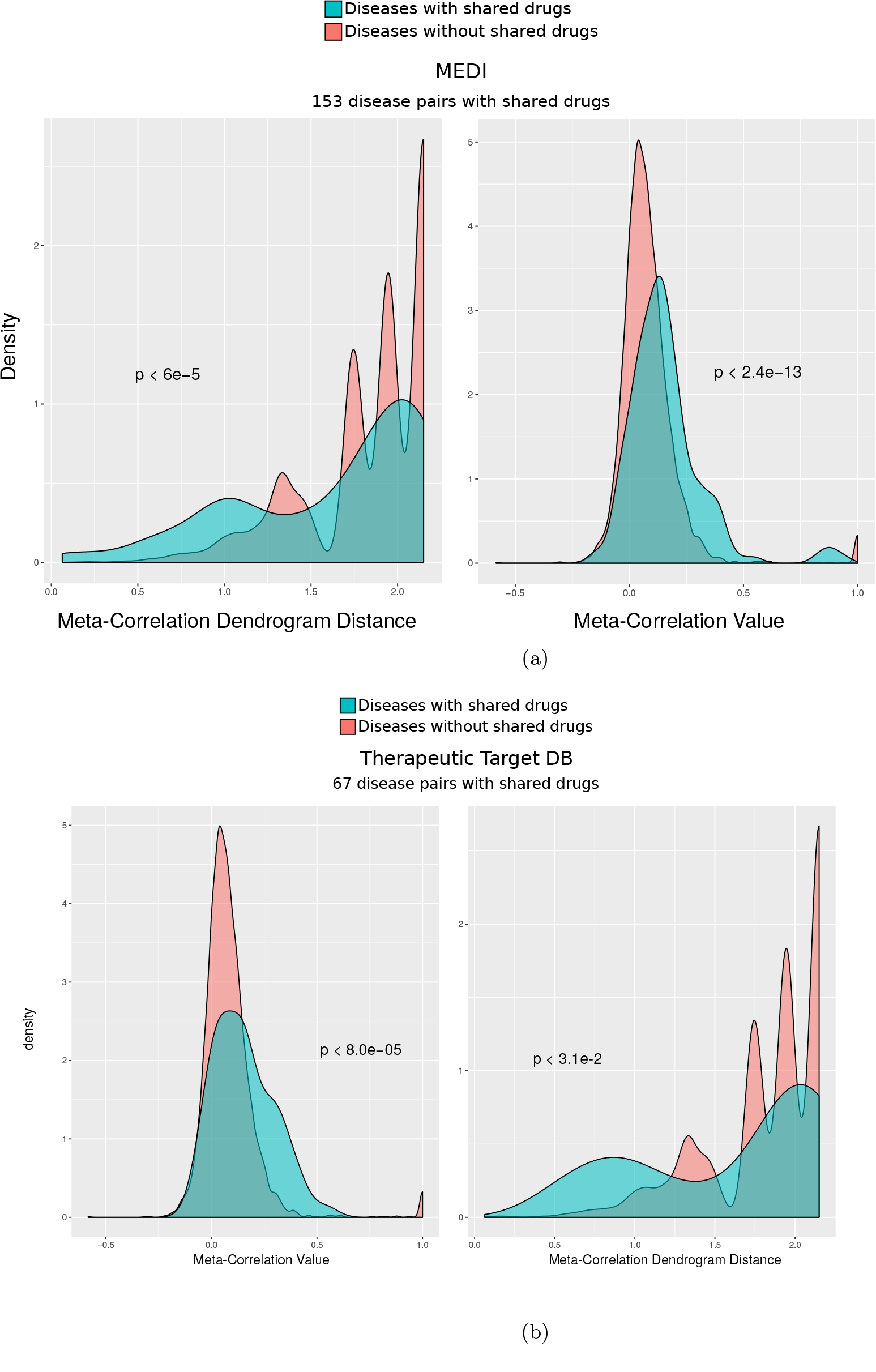
Disease pairs which share drug treatments are enriched in integrated molecular and clinical correlation analysis. We identified pairs of diseases for which the same drug is indicated in MEDI^59^ and Therapeutics Target DB.^82^ We compared these disease pairs to the diseases which did not share indications in terms of the distance in the integrated molecular and clinical correlation dendrogram and the integrated correlation values. Across all comparisons, the diseases which shared drugs were significantly more similar than the background population by the Wilcox rank-sum test (p-values indicated on plots).

**Figure S5:**
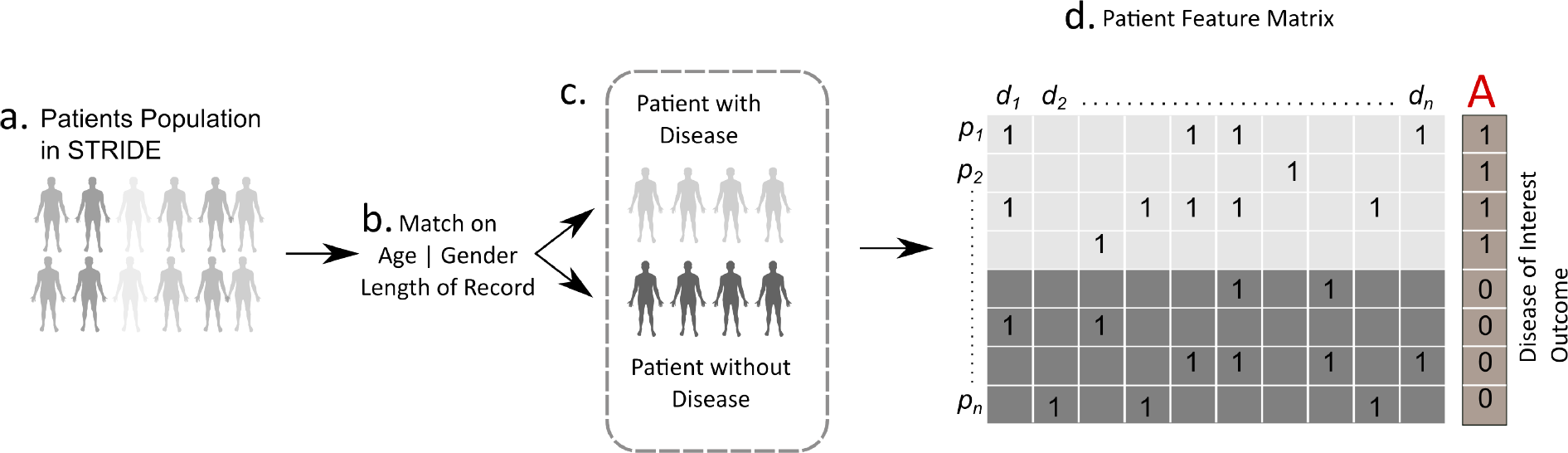
Procedure to compute clinical effect sizes. a) Patient population in STRIDE b) Patient with disease matched with other patients in the population based on age, gender and length of record to find similar patients without disease c) A disease cohort comprising patients with disease and matched patients without disease d) Patient feature matrix of a given disease cohort where rows are the patients in cohort and column are other diseases coded in the record of a given patient.

**Figure S6:**
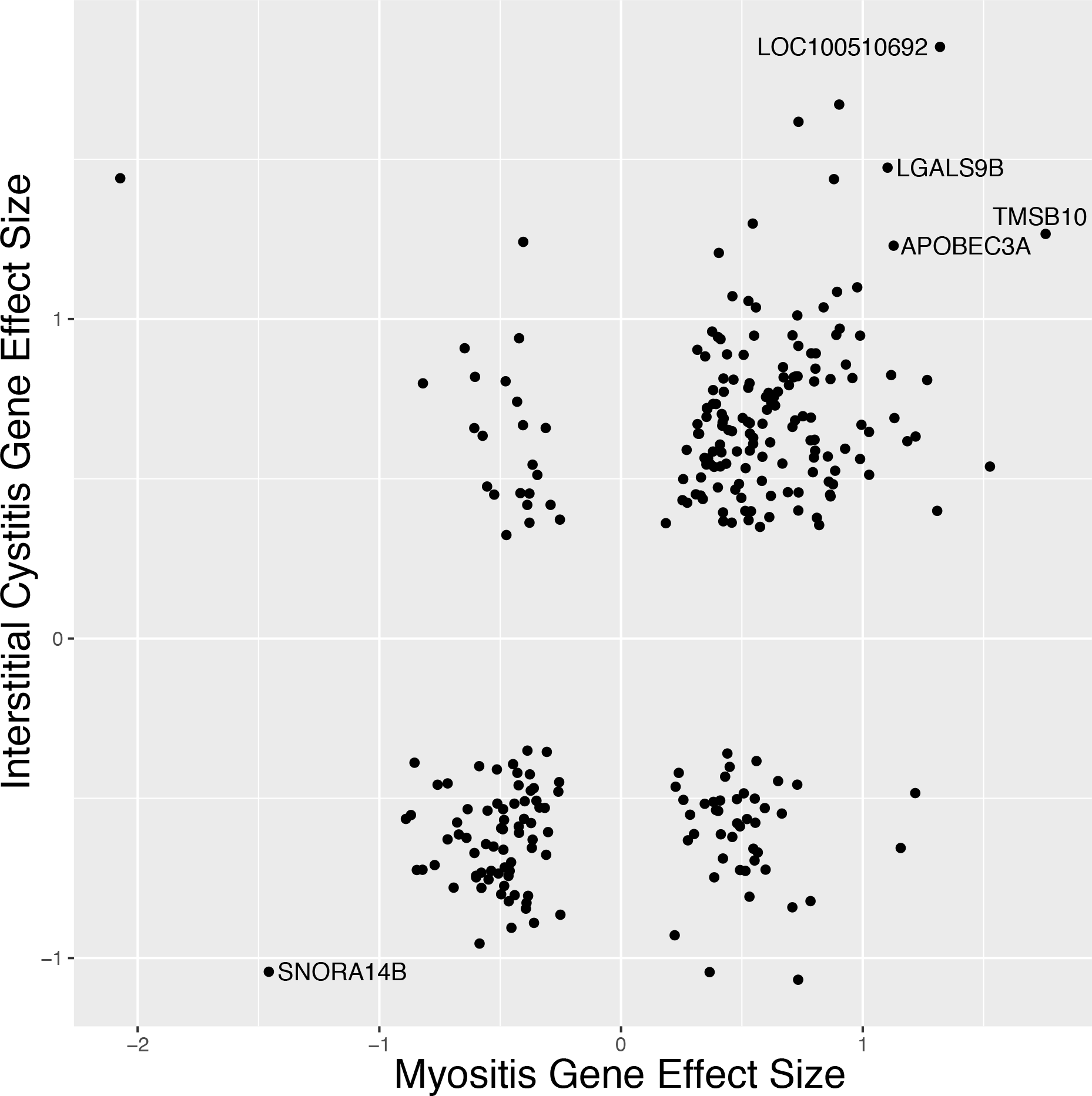
Genes that were significant at an FDR of less than 5% in both myositis and interstitial cystitis. Text labels identify the genes that were most consistently expressed in both diseases.

## References

1 S. M. Leach, H. Tipney, W. Feng, W. A. Baumgartner Jr., P. Kasliwal, R. P. Schuyler, T. Williams, R. A. Spritz, and L. Hunter, “Biomedical Discovery Acceleration, with Applications to Craniofacial Development,” PLoS Comput Biol, vol. 5, no. 3, pp. 1–19, 2009. [Online]. Available: http://dx.doi.org/10.1371%2Fjournal.pcbi.1000215

2 R. B. Altman, M. Bada, X. J. Chai, M. W. Carillo, R. O. Chen, and N. F. Abernethy, “RiboWeb: An ontology-based system for collaborative molecular biology,” IEEE Intelligent Systems, vol. 14, no. 5, pp. 68–76, 1999.

3 K.-I. Goh, M. E. Cusick, D. Valle, B. Childs, M. Vidal, and A.-L. Barabási, “The human disease network.” Proceedings of the National Academy of Sciences of the United States of America, vol. 104, no. 21, pp. 8685–8690, 2007. [Online]. Available: http://www.pubmedcentral.nih.gov/articlerender.fcgi?artid=1885563&tool=pmcentrez&rendertype=abstract

4 B. Bulik-Sullivan, H. K. Finucane, V. Anttila, A. Gusev, F. R. Day, P.-R. Loh, L. Duncan, J. R. B. Perry, N. Patterson, E. B. Robinson, M. J. Daly, A. L. Price, and B. M. Neale, “An atlas of genetic correlations across human diseases and traits.” Nature genetics, vol. advance on, 9 2015. [Online]. Available: http://dx.doi.org/10.1038/ng.3406

5 J. Menche, A. Sharma, M. Kitsak, S. D. Ghiassian, M. Vidal, J. Loscalzo, and A.-L. Barabasi, “Uncovering disease-disease relationships through the incomplete interactome,” Science, vol. 347, no. 6224, pp. 1257601–, 2 2015. [Online]. Available: http://www.sciencemag.org/content/347/6224/1257601.full

6 K. Lage, E. O. Karlberg, Z. M. Størling, P. I. Olason, A. G. Pedersen, O. Rigina, A. M. Hinsby, Z. Tümer, F. Pociot, N. Tommerup, Y. Moreau, and S. Brunak, “A human phenome-interactome network of protein complexes implicated in genetic disorders.” Nature biotechnology, vol. 25, no. 3, pp. 309–16, 3 2007. [Online]. Available: http://dx.doi.org/10.1038/nbt1295

7 S. C. Bagley, M. Sirota, R. Chen, A. J. Butte, and R. B. Altman, “Constraints on Biological Mechanism from Disease Comorbidity Using Electronic Medical Records and Database of Genetic Variants.” PLoS computational biology, vol. 12, no. 4, p. e1004885, 4 2016. [Online]. Available: http://journals.plos.org/ploscompbiol/article?id=10.1371%2Fjournal.pcbi.1004885

8 J. K. Pickrell, T. Berisa, J. Z. Liu, L. Ségurel, J. Y. Tung, and D. A. Hinds, “Detection and interpretation of shared genetic influences on 42 human traits,” Nature Genetics, vol. advance on, 5 2016. [Online]. Available: http://dx.doi.org/10.1038/ng.3570

9 D. Gligorijevic, J. Stojanovic, N. Djuric, V. Radosavljevic, M. Grbovic, R. J. Kulathinal, and Z. Obradovic, “Large-Scale Discovery of Disease-Disease and Disease-Gene Associations.” Scientific reports, vol. 6, p. 32404, 2016. [Online]. Available: http://www.ncbi.nlm.nih.gov/pubmed/27578529

10 D. R. Blair, C. S. Lyttle, J. M. Mortensen, C. F. Bearden, A. B. Jensen, H. Khiabanian, R. Melamed, R. Rabadan, E. V. Bernstam, S. Brunak, L. J. Jensen, D. Nicolae, N. H. Shah, R. L. Grossman, N. J. Cox, K. P. White, and A. Rzhetsky, “A nondegenerate code of deleterious variants in Mendelian loci contributes to complex disease risk.” Cell, vol. 155, no. 1, pp. 70–80, 9 2013. [Online]. Available: http://www.sciencedirect.com/science/article/pii/S0092867413010246

11 A. Verma, A. O. Basile, Y. Bradford, H. Kuivaniemi, G. Tromp, D. Carey, G. S. Gerhard, J. E. Crowe, M. D. Ritchie, and S. A. Pendergrass, “Phenome-Wide Association Study to Explore Relationships between Immune System Related Genetic Loci and Complex Traits and Diseases,” PLOS ONE, vol. 11, no. 8, p. e0160573, 8 2016. [Online]. Available: http://www.ncbi.nlm.nih.gov/pubmed/27508393http://www.pubmedcentral.nih.gov/articlerender.fcgi?artid=PMC4980020http://dx.plos.org/10.1371/journal.pone.0160573

12 D. M. Roden, “Phenome-wide association studies: a new method for functional genomics in humans,” The Journal of Physiology, 3 2017. [Online]. Available: http://www.ncbi.nlm.nih.gov/pubmed/28229460http://doi.wiley.com/10.1113/JP273122

13 A. Verma, S. S. Verma, S. A. Pendergrass, D. C. Crawford, D. R. Crosslin, H. Kuivaniemi, W. S. Bush, Y. Bradford, I. Kullo, S. J. Bielinski, R. Li, J. C. Denny, P. Peissig, S. Hebbring, M. De Andrade, M. D. Ritchie, and G. Tromp, “eMERGE Phenome-Wide Association Study (PheWAS) identifies clinical associations and pleiotropy for stop-gain variants,” BMC Medical Genomics, vol. 9, no. S1, p. 32, 8 2016. [Online]. Available: http://www.ncbi.nlm.nih.gov/pubmed/27535653http://www.pubmedcentral.nih.gov/articlerender.fcgi?artid=PMC4989894http://bmcmedgenomics.biomedcentral.com/articles/10.1186/s12920-016-0191-8

14 N. H. Shah, “Mining the ultimate phenome repository.” Nature biotechnology, vol. 31, no. 12, pp. 1095–7, 12 2013. [Online]. Available: http://www.ncbi.nlm.nih.gov/pubmed/24316646http://www.pubmedcentral.nih.gov/articlerender.fcgi?artid=PMC4036679

15 D. A. Davis and N. V. Chawla, “Exploring and Exploiting Disease Interactions from Multi-Relational Gene and Phenotype Networks,” PLoS ONE, vol. 6, no. 7, p. e22670, 7 2011. [Online]. Available: http://dx.plos.org/10.1371/journal.pone.0022670

16 S. Suthram, J. T. Dudley, A. P. Chiang, R. Chen, T. J. Hastie, and A. J. Butte, “Network-based elucidation of human disease similarities reveals common functional modules enriched for pluripotent drug targets.” PLoS computational biology, vol. 6, no. 2, p. e1000662, 2 2010. [Online]. Available: http://journals.plos.org/ploscompbiol/article?id=10.1371/journal.pcbi.1000662

17 D. R. Rhodes, J. Yu, K. Shanker, N. Deshpande, R. Varambally, D. Ghosh, T. Barrette, A. Pandey, and A. M. Chinnaiyan, “ONCOMINE: a cancer microarray database and integrated data-mining platform.” Neoplasia, vol. 6, no. 1, pp. 1–6, 1 2004. [Online]. Available: http://www.pubmedcentral.nih.gov/articlerender.fcgi?artid=1635162&tool=pmcentrez&rendertype=abstract

18 J. Chen, J. Cerise, A. Jabbari, R. Clynes, and A. Christiano, “Master Regulators of Infiltrate Recruitment in Autoimmune Disease Identified through Network-Based Molecular Deconvolution,” Cell Systems, vol. 1, no. 5, pp. 326–337, 11 2015. [Online]. Available: http://www.ncbi.nlm.nih.gov/pubmed/26665180http://www.pubmedcentral.nih.gov/articlerender.fcgi?artid=PMC4670983http://linkinghub.elsevier.com/retrieve/pii/S2405471215001878

19 T. Tuller, S. Atar, E. Ruppin, M. Gurevich, and A. Achiron, “Common and specific signatures of gene expression and proteinâĂŞprotein interactions in autoimmune diseases,” Genes and Immunity, vol. 14, no. 2, pp. 67–82, 3 2013. [Online]. Available: http://www.ncbi.nlm.nih.gov/pubmed/23190644http://www.nature.com/doifinder/10.1038/gene.2012.55

20 T. E. Sweeney, L. Braviak, C. M. Tato, and P. Khatri, “Genome-wide expression for diagnosis of pulmonary tuberculosis: a multicohort analysis,” The Lancet Respiratory Medicine, vol. 4, no. 3, pp. 213–224, 2016.

21 C. A. Hidalgo, N. Blumm, A.-L. Barabási, and N. A. Christakis, “A Dynamic Network Approach for the Study of Human Phenotypes,” PLoS Computational Biology, vol. 5, no. 4, p. e1000353, 4 2009. [Online]. Available: http://dx.plos.org/10.1371/journal.pcbi.1000353

22 F. S. Roque, P. B. Jensen, H. Schmock, M. Dalgaard, M. Andreatta, T. Hansen, K. Søeby, S. Bredkjær, A. Juul, T. Werge, L. J. Jensen, and S. Brunak, “Using Electronic Patient Records to Discover Disease Correlations and Stratify Patient Cohorts,” PLoS Computational Biology, vol. 7, no. 8, p. e1002141, 8 2011. [Online]. Available: http://dx.plos.org/10.1371/journal.pcbi.1002141

23 E. Capobianco and P. LioâĂŹ, “Comorbidity: a multidimensional approach,” Trends in Molecular Medicine, vol. 19, no. 9, pp. 515–521, 2013.

24 A. B. Jensen, P. L. Moseley, T. I. Oprea, S. G. Ellesøe, R. Eriksson, H. Schmock, P. B. Jensen, L. J. Jensen, S. Brunak, O. Camilo, L. B. Goldstein, J. Finkelstein, E. Cha, S. M. Scharf, J. M. Teno, S. Weitzen, M. L. Fenell, V. Mor, F. E. M. Murtagh, E. Murphy, N. S. Sheerin, F. E. M. Murtagh, N. S. Sheerin, J. Addington-Hall, I. J. Higginson, S. A. Murray, M. Kendall, K. Boyd, A. Sheikh, H. Petri, D. Maldonato, N. J. Robinson, D. R. Blair, C. A. Hidalgo, N. Blumm, A.-L. Barabási, N. A. Christakis, L. L. Chen, N. Blumm, N. A. Christakis, A.-L. Barabási, T. S. Deisboeck, H. Tanushi, H. Dalianis, G. H. Nilsson, S. M. Curkendall, S. Sidney, A. C. Salisbury, K. J. Reid, J. A. Spertus, S. Suissa, S. DellâĂŹAniello, P. Ernst, S. E. Moss, R. Klein, B. E. K. Klein, T. Y. Wong, E. M. Kohner, D. S. Freedman, D. F. Williamson, E. W. Gunter, T. Byers, A. Kelkar, A. Kuo, W. H. Frishman, Q. Yang, B. M. Farr, C. L. Bartlett, J. Wadsworth, D. L. Miller, and T. S. Ingebrigtsen, “Temporal disease trajectories condensed from population-wide registry data covering 6.2 million patients,” Nature Communications, vol. 5, pp. 1769–1775, 6 2014. [Online]. Available: http://www.nature.com/doifinder/10.1038/ncomms5022

25 E. Capobianco and P. Liò, “Comorbidity networks: beyond disease correlations,” Journal of Complex Networks, vol. 3, no. 3, pp. 319–332, 9 2015. [Online]. Available: http://comnet.oxfordjournals.org/lookup/doi/10.1093/comnet/cnu048

26 J. X. Hu, C. E. Thomas, and S. Brunak, “Network biology concepts in complex disease comorbidities,” Nature Reviews Genetics, vol. 17, 8 2016. [Online]. Available: http://www.nature.com/doifinder/10.1038/nrg.2016.87

27 L. Li, W.-Y. Cheng, B. S. Glicksberg, O. Gottesman, R. Tamler, R. Chen, E. P. Bottinger, and J. T. Dudley, “Identification of type 2 diabetes subgroups through topological analysis of patient similarity,” Science Translational Medicine, vol. 7, no. 311, 2015. [Online]. Available: http://stm.sciencemag.org.ezproxy.stanford.edu/content/7/311/311ra174.full

28 A. Rzhetsky, D. Wajngurt, N. Park, and T. Zheng, “Probing genetic overlap among complex human phenotypes,” Proceedings of the National Academy of Sciences, vol. 104, no. 28, pp. 11 694–11 699, 7 2007. [Online]. Available: http://www.ncbi.nlm.nih.gov/pubmed/17609372http://www.pubmedcentral.nih.gov/articlerender.fcgi?artid=PMC1906727http://www.pnas.org/cgi/doi/10.1073/pnas.0704820104

29 T. T. Ashburn and K. B. Thor, “Drug repositioning: identifying and developing new uses for existing drugs,” Nature Reviews Drug Discovery, vol. 3, no. 8, pp. 673–683, 8 2004. [Online]. Available: http://www.nature.com/doifinder/10.1038/nrd1468

30 D. Sardana, C. Zhu, M. Zhang, R. C. Gudivada, L. Yang, and A. G. Jegga, “Drug repositioning for orphan diseases.” Briefings in bioinformatics, vol. 12, no. 4, pp. 346–56, 7 2011. [Online]. Available: http://www.ncbi.nlm.nih.gov/pubmed/21504985

31 P. B. Jensen, L. J. Jensen, and S. Brunak, “Mining electronic health records: towards better research applications and clinical care,” Nature Reviews Genetics, vol. 13, no. 6, pp. 395–405, 5 2012. [Online]. Available: http://www.ncbi.nlm.nih.gov/pubmed/22549152http://www.nature.com/doifinder/10.1038/nrg3208

32 S. J. Shah, D. H. Katz, S. Selvaraj, M. A. Burke, C. W. Yancy, M. Gheorghiade, R. O. Bonow, C.-C. Huang, and R. C. Deo, “Phenomapping for Novel Classification of Heart Failure With Preserved Ejection Fraction,” Circulation, vol. 131, no. 3, pp. 269–279, 1 2015. [Online]. Available: http://www.ncbi.nlm.nih.gov/pubmed/25398313http://www.pubmedcentral.nih.gov/articlerender.fcgi?artid=PMC4302027http://circ.ahajournals.org/cgi/doi/10.1161/CIRCULATIONAHA.114.010637

33 Y. S. Low, A. C. Daugherty, E. A. Schroeder, W. Chen, T. Seto, S. Weber, M. Lim, T. Hastie, M. Mathur, M. Desai, C. Farrington, A. A. Radin, M. Sirota, P. Kenkare, C. A. Thompson, P. P. Yu, S. L. Gomez, G. W. Sledge, A. W. Kurian, and N. H. Shah, “Synergistic drug combinations from electronic health records and gene expression,” Journal of the American Medical Informatics Association, vol. 24, no. 3, p. ocw161, 12 2016. [Online]. Available: http://www.ncbi.nlm.nih.gov/pubmed/27940607https://academic.oup.com/jamia/article-lookup/doi/10.1093/jamia/ocw161

34 H. Xu, M. C. Aldrich, Q. Chen, H. Liu, N. B. Peterson, Q. Dai, M. Levy, A. Shah, X. Han, X. Ruan, M. Jiang, Y. Li, J. S. Julien, J. Warner, C. Friedman, D. M. Roden, and J. C. Denny, “Validating drug repurposing signals using electronic health records: a case study of metformin associated with reduced cancer mortality,” Journal of the American Medical Informatics Association, vol. 22, no. 1, pp. 179–91, 7 2014. [Online]. Available: http://www.ncbi.nlm.nih.gov/pubmed/25053577http://www.pubmedcentral.nih.gov/articlerender.fcgi?artid=PMC4433365https://academic.oup.com/jamia/article-lookup/doi/10.1136/amiajnl-2014-002649

35 L. M. Schriml, C. Arze, S. Nadendla, Y. W. W. Chang, M. Mazaitis, V. Felix, G. Feng, and W. A. Kibbe, “Disease ontology: A backbone for disease semantic integration,” Nucleic Acids Research, vol. 40, no. D1, 2012.

36 “International Classification of Diseases, 10th Revision, Clinical Modification (ICD-10-CM),” National Center for Health Statistic, vol. 1, 2015.

37 L. Cheng, Y. Jiang, Z. Wang, H. Shi, J. Sun, H. Yang, S. Zhang, Y. Hu, and M. Zhou, “DisSim: an online system for exploring significant similar diseases and exhibiting potential therapeutic drug,” Scientific Reports, vol. 6, no. 1, p. 30024, 9 2016. [Online]. Available: http://www.ncbi.nlm.nih.gov/pubmed/27457921http://www.pubmedcentral.nih.gov/articlerender.fcgi?artid=PMC4960572http://www.nature.com/articles/srep30024

38 R. Pivovarov and N. Elhadad, “A hybrid knowledge-based and data-driven approach to identifying semantically similar concepts,” Journal of Biomedical Informatics, vol. 45, no. 3, pp. 471–481, 6 2012. [Online]. Available: http://www.ncbi.nlm.nih.gov/pubmed/22289420http://www.pubmedcentral.nih.gov/articlerender.fcgi?artid=PMC3345313http://linkinghub.elsevier.com/retrieve/pii/S1532046412000032

39 J. Li, B. Gong, X. Chen, T. Liu, C. Wu, F. Zhang, C. Li, X. Li, S. Rao, and X. Li, “DOSim: An R package for similarity between diseases based on Disease Ontology,” BMC Bioinformatics, vol. 12, no. 1, p. 266, 6 2011. [Online]. Available: http://www.ncbi.nlm.nih.gov/pubmed/21714896http://www.pubmedcentral.nih.gov/articlerender.fcgi?artid=PMC3150296http://bmcbioinformatics.biomedcentral.com/articles/10.1186/1471-2105-12-266

40 W. A. Haynes, F. Vallania, C. Liu, E. Bongen, A. Tomczak, M. Andres-Terrè, S. Lofgren, A. Tam, C. A. Deisseroth, M. D. Li, T. E. Sweeney, and P. Khatri, “Empowering Multi-Cohort Gene Expression Analysis to Increase Reproducibility,” Pac Symp Biocomput, vol. Web, 2017. [Online]. Available: http://biorxiv.org/content/early/2016/08/25/071514

41 T. E. Sweeney, W. A. Haynes, F. Vallania, J. P. Ioannidis, and P. Khatri, “Methods to increase reproducibility in differential gene expression via meta-analysis.” Nucleic acids research, vol. Web, p. gkw797, 9 2016. [Online]. Available: http://www.ncbi.nlm.nih.gov/pubmed/27634930

42 W. Haynes, A. Tomczak, and P. Khatri, “Gene annotation bias impedes biomedical research,” Pacific Symposium on Biocomputing, 2017. [Online]. Available: http://biorxiv.org/content/early/2017/05/02/133108

43 P. Khatri, S. Roedder, N. Kimura, K. De Vusser, A. A. Morgan, Y. Gong, M. P. Fischbein, R. C. Robbins, M. Naesens, A. J. Butte, and M. M. Sarwal, “A common rejection module (CRM) for acute rejection across multiple organs identifies novel therapeutics for organ transplantation.” The Journal of experimental medicine, vol. 210, no. 11, pp. 2205–21, 10 2013. [Online]. Available: http://jem.rupress.org/content/210/11/2205.full

44 R. Chen, P. Khatri, P. K. Mazur, M. Polin, Y. Zheng, D. Vaka, C. D. Hoang, J. Shrager, Y. Xu, S. Vicent, A. J. Butte, and E. A. Sweet-Cordero, “A meta-analysis of lung cancer gene expression identifies PTK7 as a survival gene in lung adenocarcinoma,” Cancer Research, vol. 74, no. 10, pp. 2892–2902, 5 2014. [Online]. Available: http://www.ncbi.nlm.nih.gov/pubmed/2465423

45 T. E. Sweeney, A. Shidham, H. R. Wong, and P. Khatri, “A comprehensive time-course-based multicohort analysis of sepsis and sterile inflammation reveals a robust diagnostic gene set.” Science Translational Medicine, vol. 7, no. 287, p. 287ra71, 5 2015. [Online]. Available: http://stm.sciencemag.org/content/7/287/287ra71.abstract

46 M. Andres-Terre, H. McGuire, Y. Pouliot, E. Bongen, T. Sweeney, C. Tato, and P. Khatri, “Integrated, Multi-cohort Analysis Identifies Conserved Transcriptional Signatures across Multiple Respiratory Viruses,” Immunity, vol. 43, no. 6, pp. 1199–1211, 12 2015. [Online]. Available: http://www.cell.com/article/S1074761315004550/fulltext

47 M. D. Li, T. C. Burns, A. A. Morgan, and P. Khatri, “Integrated multi-cohort transcriptional meta-analysis of neurodegenerative diseases.” Acta neuropathologica communications, vol. 2, p. 93, 1 2014. [Online]. Available: http://www.pubmedcentral.nih.gov/articlerender.fcgi?artid=4167139&tool=pmcentrez&rendertype=abstract

48 P. K. Mazur, N. Reynoird, P. Khatri, P. W. T. C. Jansen, A. W. Wilkinson, S. Liu, O. Barbash, G. S. Van Aller, M. Huddleston, D. Dhanak, P. J. Tummino, R. G. Kruger, B. A. Garcia, A. J. Butte, M. Vermeulen, J. Sage, and O. Gozani, “SMYD3 links lysine methylation of MAP3K2 to Ras-driven cancer.” Nature, vol. advance on, 5 2014. [Online]. Available: www.nature.com/articles/nature13320

49 P. K. Mazur, A. Herner, S. S. Mello, M. Wirth, S. Hausmann, F. J. Sánchez-Rivera, S. M. Lofgren, T. Kuschma, S. A. Hahn, D. Vangala, M. Trajkovic-Arsic, A. Gupta, I. Heid, P. B. Noël, R. Braren, M. Erkan, J. Kleeff, B. Sipos, L. C. Sayles, M. Heikenwalder, E. Heßmann, V. Ellenrieder, I. Esposito, T. Jacks, J. E. Bradner, P. Khatri, E. A. Sweet-Cordero, L. D. Attardi, R. M. Schmid, G. Schneider, J. Sage, and J. T. Siveke, “Combined inhibition of BET family proteins and histone deacetylases as a potential epigenetics-based therapy for pancreatic ductal adenocarcinoma,” Nature Medicine, vol. 21, no. 10, pp. 1163–1171, 9 2015. [Online]. Available: http://www.nature.com/doifinder/10.1038/nm.3952

50 T. E. Sweeney, H. R. Wong, and P. Khatri, “Robust classification of bacterial and viral infections via integrated host gene expression diagnostics.” Science translational medicine, vol. 8, no. 346, p. 346ra91, 7 2016. [Online]. Available: http://www.ncbi.nlm.nih.gov/pubmed/27384347

51 H.-C. S. P. HIPC-CHI Signatures Project Team and H.-I. HIPC-I Consortium, “Multicohort analysis reveals baseline transcriptional predictors of influenza vaccination responses.” Science immunology, vol. 2, no. 14, p. eaal4656, 8 2017. [Online]. Available: http://www.ncbi.nlm.nih.gov/pubmed/28842433

52 H. J. Lowe, T. A. Ferris, P. M. Hernandez, and S. C. Weber, “STRIDE–An integrated standards-based translational research informatics platform.” AMIA … Annual Symposium proceedings/AMIA Symposium. AMIA Symposium, vol. 2009, pp. 391–5, 2009. [Online]. Available: http://www.ncbi.nlm.nih.gov/pubmed/20351886http://www.pubmedcentral.nih.gov/articlerender.fcgi?artid=PMC2815452

53 J. A. James, B. R. Neas, K. L. Moser, T. Hall, G. R. Bruner, A. L. Sestak, and J. B. Harley, “Systemic lupus erythematosus in adults is associated with previous Epstein-Barr virus exposure,” Arthritis and Rheumatism, vol. 44, no. 5, pp. 1122–1126, 2001.

54 H. Tadema, P. Heeringa, and C. G. M. Kallenberg, “Bacterial infections in Wegener’s granulomatosis: mechanisms potentially involved in autoimmune pathogenesis.” Current opinion in rheumatology, vol. 23, pp. 366–371, 2011.

55 O. M. Kon and R. M. duBois, “Mycobacteria and sarcoidosis,” Thorax, vol. 52, pp. S47–S51, 1997.

56 C. Pagnoux, P. Cohen, and L. Guillevin, “Vasculitides secondary to infections.” Clinical and experimental rheumatology, vol. 24, no. 2 Suppl 41, pp. 71–81, 2006. [Online]. Available: http://www.ncbi.nlm.nih.gov/pubmed/16859600

57 K. M. W. Tauil, E. Gaio, C. A. Melo-Silva, R. S. Carvalho, and V. M. Amado, “Pulmonary arterial hypertension and sepsis: prothrombotic profile and inflammation can changes pulmonary mechanics?” Medical hypotheses, vol. 83, no. 3, pp. 290–1, 9 2014. [Online]. Available: http://www.ncbi.nlm.nih.gov/pubmed/24957506

58 S. Janda, B. S. Quon, and J. Swiston, “HIV and pulmonary arterial hypertension: a systematic review.” HIV medicine, vol. 11, no. 10, pp. 620–34, 11 2010. [Online]. Available: http://www.ncbi.nlm.nih.gov/pubmed/20408888

59 W.-Q. Wei, R. M. Cronin, H. Xu, T. A. Lasko, L. Bastarache, and J. C. Denny, “Development and evaluation of an ensemble resource linking medications to their indications.” Journal of the American Medical Informatics Association: JAMIA, vol. 20, no. 5, pp. 954–61, 2013. [Online]. Available: http://www.ncbi.nlm.nih.gov/pubmed/23576672http://www.pubmedcentral.nih.gov/articlerender.fcgi?artid=PMC3756263

60 M. Ashburner, C. A. Ball, J. A. Blake, D. Botstein, H. Butler, J. M. Cherry, A. P. Davis, K. Dolinski, S. S. Dwight, J. T. Eppig, M. A. Harris, D. P. Hill, L. Issel-Tarver, A. Kasarskis, S. Lewis, J. C. Matese, J. E. Richardson, M. Ringwald, G. M. Rubin, and G. Sherlock, “Gene ontology: tool for the unification of biology. The Gene Ontology Consortium.” Nature genetics, vol. 25, no. 1, pp. 25–9, 5 2000. [Online]. Available: http://dx.doi.org/10.1038/75556

61 W. A. Haynes, A. Tomczak, and P. Khatri, “Gene annotation bias impedes biomedical research,” Scientific Reports, vol. 8, no. 1, p. 1362, 12 2018. [Online]. Available: http://www.nature.com/articles/s41598-018-19333-x

62 H. Mi, A. Muruganujan, and P. D. Thomas, “PANTHER in 2013: modeling the evolution of gene function, and other gene attributes, in the context of phylogenetic trees.” Nucleic acids research, vol. 41, no. Database issue, pp. 377–86, 1 2013. [Online]. Available: http://www.pubmedcentral.nih.gov/articlerender.fcgi?artid=3531194&tool=pmcentrez&rendertype=abstract

63 J. T. Dudley, R. Tibshirani, T. Deshpande, and A. J. Butte, “Disease signatures are robust across tissues and experiments.” Molecular systems biology, vol. 5, p. 307, 2009. [Online]. Available: http://www.ncbi.nlm.nih.gov/pubmed/19756046http://www.pubmedcentral.nih.gov/articlerender.fcgi?artid=PMC2758720

64 A. Brazma, H. Parkinson, U. Sarkans, M. Shojatalab, J. Vilo, N. Abeygunawardena, E. Holloway, M. Kapushesky, P. Kemmeren, G. G. Lara, A. Oezcimen, P. Rocca-Serra, and S.-A. Sansone, “ArrayExpress–a public repository for microarray gene expression data at the EBI.” Nucleic Acids Research, vol. 31, no. 1, pp. 68–71, 1 2003. [Online]. Available: http://www.pubmedcentral.nih.gov/articlerender.fcgi?artid=165538&tool=pmcentrez&rendertype=abstract

65 R. Edgar, “Gene Expression Omnibus: NCBI gene expression and hybridization array data repository,” Nucleic Acids Research, vol. 30, no. 1, pp. 207–210, 1 2002. [Online]. Available: http://nar.oxfordjournals.org/content/30/1/207.short

66 W. Yu, M. Clyne, M. J. Khoury, and M. Gwinn, “Phenopedia and Genopedia: disease-centered and gene-centered views of the evolving knowledge of human genetic associations,” Bioinformatics, vol. 26, no. 1, pp. 145–146, 1 2010. [Online]. Available: http://bioinformatics.oxfordjournals.org/cgi/doi/10.1093/bioinformatics/btp618

67 Maggie Lam, “PubPular: Identifying the focus of biomedical research.” [Online]. Available: https://pubpular.shinyapps.io/PubPular/

68 M. P. Y. Lam, V. Venkatraman, Y. Xing, E. Lau, Q. Cao, D. C. M. Ng, A. I. Su, J. Ge, J. E. Van Eyk, and P. Ping, “Data-Driven Approach To Determine Popular Proteins for Targeted Proteomics Translation of Six Organ Systems.” Journal of proteome research, vol. Web, 7 2016. [Online]. Available: http://www.ncbi.nlm.nih.gov/pubmed/27356587

69 W. Yu, A. Yesupriya, A. Wulf, L. A. Hindorff, N. Dowling, M. J. Khoury, and M. Gwinn, “GWAS Integrator: a bioinformatics tool to explore human genetic associations reported in published genome-wide association studies,” European Journal of Human Genetics, vol. 19, no. 10, pp. 1095–1099, 10 2011. [Online]. Available: http://www.nature.com/doifinder/10.1038/ejhg.2011.91

70 D. Welter, J. MacArthur, J. Morales, T. Burdett, P. Hall, H. Junkins, A. Klemm, P. Flicek, T. Manolio, L. Hindorff, and H. Parkinson, “The NHGRI GWAS Catalog, a curated resource of SNP-trait associations.” Nucleic acids research, vol. 42, no. Database issue, pp. 1001–6, 1 2014. [Online]. Available: http://www.ncbi.nlm.nih.gov/pubmed/24316577http://www.pubmedcentral.nih.gov/articlerender.fcgi?artid=PMC3965119

71 F. FitzHenry, F. S. Resnic, S. L. Robbins, J. Denton, L. Nookala, D. Meeker, L. Ohno-Machado, and M. E. Matheny, “Creating a Common Data Model for Comparative Effectiveness with the Observational Medical Outcomes Partnership.” Applied clinical informatics, vol. 6, no. 3, pp. 536–547, 2015.

72 O. Bodenreider, “The Unified Medical Language System (UMLS): integrating biomedical terminology.” Nucleic acids research, vol. 32, no. Database issue, pp. 267–70, 1 2004. [Online]. Available: http://www.ncbi.nlm.nih.gov/pubmed/14681409http://www.pubmedcentral.nih.gov/articlerender.fcgi?artid=PMC308795

73 J. S. Sekhon, “Multivariate and Propensity Score Matching Software with Automated Balance Optimization: The Matching Package for R,” Journal of Statistical Software, vol. 55, no. 2, pp. 1–52, 2011.

74 J. Friedman, T. Hastie, and R. Tibshirani, “Regularization Paths for Generalized Linear Models via Coordinate Descent.” Journal of Statistical Software, vol. 33, no. 1, pp. 1–22, 2010. [Online]. Available: http://www.pubmedcentral.nih.gov/articlerender.fcgi?artid=2929880&tool=pmcentrez&rendertype=abstract

75 M. Borenstein, L. V. Hedges, J. P. T. Higgins, and H. R. Rothstein, Introduction to Meta-Analysis. 1. National Institutes of Health. FINAL NIH STATEMENT ON SHARING RESEARCH DATA. (2003). Available at: https://grants.nih.gov/grants/guide/notice-files/NOT-OD-03-032.html. (Accessed: 13th January 2018) 2. Nousari, H. C. & Anhalt, G. J. Bullous skin disease, 2009.

76 J. H. Ward, “Hierarchical grouping to optimize an objective function,” pp. 236–244, 1963.

77 F. Murtagh and P. Legendre, “Ward’s Hierarchical Agglomerative Clustering Method: Which Algorithms Implement Ward’s Criterion?” Journal of Classification, vol. 31, no. 3, pp. 274–295, 2014.

78 R. Suzuki and H. Shimodaira, “Pvclust: An R package for assessing the uncertainty in hierarchical clustering,” Bioinformatics, vol. 22, no. 12, pp. 1540–1542, 2006.

79 Z. Gu, L. Gu, R. Eils, M. Schlesner, and B. Brors, “Circlize implements and enhances circular visualization in R,” Bioinformatics, vol. 30, no. 19, pp. 2811–2812, 2014.

80 R. R. Sokal and F. J. Rohlf, “The Comparison of Dendrograms by Objective Methods,” Taxon, vol. 11, no. 2, pp. 33–40, 1962.

81 T. Galili, “dendextend: An R package for visualizing, adjusting and comparing trees of hierarchical clustering,” Bioinformatics, vol. 31, no. 22, pp. 3718–3720, 2015.

82 F. Zhu, Z. Shi, C. Qin, L. Tao, X. Liu, F. Xu, L. Zhang, Y. Song, X. Liu, J. Zhang, B. Han, P. Zhang, and Y. Chen, “Therapeutic target database update 2012: a resource for facilitating target-oriented drug discovery.” Nucleic acids research, vol. 40, no. Database issue, pp. 1128–36, 1 2012. [Online]. Available: http://www.ncbi.nlm.nih.gov/pubmed/21948793http://www.pubmedcentral.nih.gov/articlerender.fcgi?artid=PMC3245130

